# Entropy Changes in Water Networks Promote Protein Denaturation

**DOI:** 10.1101/2024.06.12.598657

**Authors:** Yichong Wang, Junlang Liu, Michael Peters, Ryoma Ishii, Dianzhuo Wang, Sourav Chowdhury, Kevin Kit Parker, Eugene I. Shakhnovich

**Affiliations:** Department of Chemistry and Chemical Biology, Harvard University; Cambridge, MA, 02138 USA; Disease Biophysics Group, John A. Paulson School of Engineering and Applied Sciences, Harvard University; Boston, MA, 02134 USA; Medical & Health Informatics Laboratories, NTT Research, Inc.; Sunnyvale, CA, 94085 USA; Medical Department of Biological Sciences, Birla Institute of Technology and Science-Pilani; Hyderabad, TN, 500078 India

## Abstract

For over a century, an explanation for how concentrated ions denature proteins has proven elusive. Here, we report a novel mechanism of protein denaturation driven by entropy changes in water networks. Experiments and simulations show that ion pairs like LiBr and LiCl localize water molecules and disrupt the water network’s structure, while others exert a more global effect without compromising network integrity. This disruption reduces the entropy penalty when proteins sequester water molecules during unfolding, resulting in a peculiar yet universal “inverse hydrophobic effect” that potentiates protein denaturation. Through successful isolation and systematic study of indirect solute effects, our findings offer a universal approach to salt induced protein denaturation and provide a unified framework for the decoding of the protein-water-solute nexus.

## Introduction

In the 19^th^ century, Franz Hofmeister discovered the ‘Hofmeister Series’ (*1, 2*), describing how various salts can either increase or decrease protein solubility in aqueous solutions. Interest in this phenomenon grew between the 1940s and 1960s, focusing on how concentrated inorganic ions affect protein stability and conformation (*3, 4*), with lithium bromide (LiBr) showing the most pronounced impact (*5*–*7*). While numerous studies observed distinct effects of various ions, the underlying mechanism of ion-induced denaturation lacked a comprehensive explanation due to limited experimental and theoretical approaches. In recent years, extensive research has explored the interactions between water molecules and ions (*8, 9*), offering new insights into the origin of hydrophobicity and its influence on protein folding (*10–12*), potentially driven by modifications in the intermolecular interactions of water (*13*). However, a unified framework for understanding the triadic relationship between proteins, water, and solutes remains elusive (*14*).

Despite the use of ion-induced denaturation for extracting proteins such as keratin and silk fibroin (*15, 16*), the gap between practical techniques and the lack of mechanistic insight limits our ability to fully control and utilize protein-based materials. The incomplete understanding of how ions contribute to this process has led to treating LiBr denaturants similarly to organic chemicals like urea or guanidine hydrochloride, which bind directly to proteins and require removal through dialysis (*17*). This complicates post-processing and restricts the implementation of widely adopted fabrication methods such as molding or fiber spinning (*18*). As a result, over ten million tons of keratin-rich waste are generated annually and typically disposed of through incineration or landfilling (*19*), regardless of their potential for applications ranging from textiles to tissue engineering scaffolds (*20, 21*).

The phenomenon of protein denaturation is governed by the balance between opposing enthalpic and entropic driving forces. The gain in protein conformational entropy drives towards unfolding, as the polypeptide chain becomes more disordered. In contrast, enthalpic interactions between amino acids (e.g., Van der Waals forces, electrostatic interactions) and the increase in entropy of water molecules released from their interactions with non-polar side chains of proteins—known as the hydrophobic effect—stabilize the folded state under native conditions. Achieving denaturation requires overcoming the entropy penalty from the loss of potential water conformations as free water molecules are sequestered by proteins upon unfolding. One possible explanation for protein denaturation in concentrated ionic solutions is the indirect solute effect, which modulates the hydrophobic effect by altering the water hydrogen bond network (*13*).

To test this hypothesis, we selected a series of inorganic ion pairs—lithium bromide (LiBr), lithium chloride (LiCl), and sodium bromide (NaBr)—as model solutes, given their similar composition but distinct reported impacts on protein structures (*3*). We systematically explored the denaturation potency of these ionic solutes by evaluating their effects on three proteins spanning different levels of structural complexity, with LiBr emerging as the strongest universal denaturant and NaBr as the least potent. Both experimental and theoretical analyses revealed that these differences stem from the ion pairs’ varying capabilities to modulate the water network rather than directly binding to the proteins. Through their local interactions with water molecules, ions like LiBr can significantly alter the structure of the water network, thereby reducing the entropy penalty associated with protein unfolding. Furthermore, this entropic denaturation mechanism offers significant advantages for extracting and regenerating keratin materials, enabling closed-loop recycling of the denaturant, simplifying the manufacturing process, and preserving the native shape-memory property.

### Universal denaturation capacity of LiBr

To systematically investigate solute effects on protein conformation, we selected three proteins representing different levels of structural complexity (Fig. 1A), along with three different ionic solutes including LiBr, LiCl, and NaBr. We found markedly distinct solute effects for different proteins and salts across various concentrations, as evidenced by aggregation of denatured protein during turbidity measurements at 405 nm (Fig. 1, B to D). The smallest protein tested, dihydrofolate reductase (DHFR) showed significant aggregation at 1 M LiBr and 5 M LiCl, but remained soluble at all NaBr concentrations (Fig. 1B, and fig. S1). Since both salt-induced precipitation and denaturation can result in protein aggregation, we used Fourier-transform infrared spectroscopy (FTIR) to further distinguish the underlying mechanisms. Starting from 2 M LiBr, we observed a gradual loss of secondary structure of DHFR such as α-helix (1650-1655 cm^-1^) and β-sheet/turns (1660-1690 cm^-1^), along with an increase in aggregated β-like random structures (1620-1630 cm^-1^) (Fig. 1E). The secondary structure loss of DHFR in 2 M LiBr was further confirmed by conformational kinetics measurement using quantitative Raman spectroscopy, where less-stable β-sheet disappeared first, followed by α-helix, with obvious accumulation of aggregated random structures (Fig. 1H, and figs. S2 and S3). Both FTIR and tryptophan fluorescence measurement confirmed DHFR unfolding starting from ∼5 M LiCl (fig. S4, A and C). In contrast, DHFR stayed well-folded for all concentrations of NaBr.

**Fig. 1.**
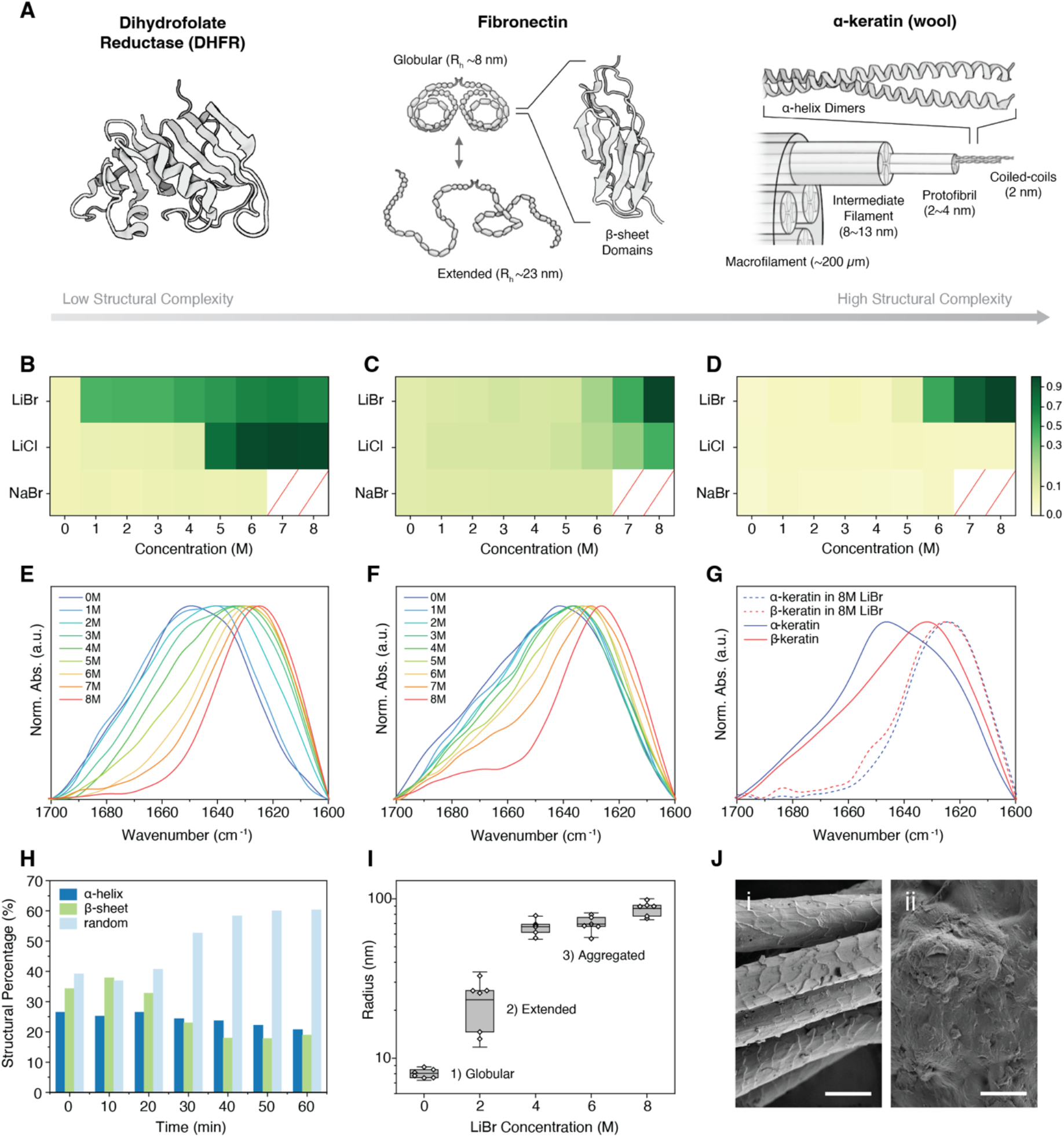
Concentrated LiBr solutions induce universal protein conformational change and cause denaturation. (**A**), Schematics of DHFR, fibronectin, and α-keratin (wool), representing three levels of structural complexity. (**B-D**) Turbidity assay of three respective proteins measured by OD405, suggesting different denaturation capabilities of LiBr, LiCl, and NaBr. Data are normalized by the highest OD405 values of respective proteins and presented as mean (n = 4). (**E-G**), FTIR of the amide I band indicate a gradual loss of secondary structure upon increasing of LiBr concentration. (**H**), Percentage change of the secondary structure in DHFR in 2 M LiBr, deconvoluted from the amide I band of Raman spectra. (**I**), DLS measurement of the hydrodynamic radius of fibronectin with increasing LiBr concentration, indicating an extension of quaternary structure followed by aggregation. Data are presented as mean ± 1.5 s.d. (n = 6). (**J**), SEM image of wool prior (i) and after (ii) denaturation with 8 M LiBr. Scale bars, 50 μm.

To elucidate whether ionic solutes can affect conformations of proteins with more complex, hierarchically ordered structures, we examined fibronectin. This protein, a critical component for cell-extracellular matrix interactions (*22*) has more recently been explored as a synthetic building material (*23–25*) because of its unique structure-function relationships. Fibronectin is comprised of two covalently linked extensible arms formed by β-sheet domains that can undergo quaternary structure changes from globular to extended state (*26*) (Fig. 1A middle). With the increase of protein structural complexity, higher concentrations of LiBr and LiCl (7 M and 8 M, respectively) were required to induce aggregation of fibronectin (Fig. 1C, and fig. S5), where secondary structure loss was also confirmed by FTIR spectra (Fig. 1F, and fig. S4B). Before denaturation, protein conformational changes were observed through dynamic light scattering (DLS) measurements of the hydrodynamic radius R_h_ (Fig. 1I), where fibronectin transitions from the globular state (R_h_ ∼8 nm) to the extended state (R_h_ ∼23 nm) at a lower concentration of LiBr around 2 M. Combining the loss of the secondary and tertiary structure observed in FTIR and the quaternary structure extension shown via DLS, we can identify sequential changes in protein structure as the concentration of LiBr increases (fig. S4D).

We also tested keratin structural transformation in LiBr solution as observed by Feughelman et al. (*3*). In contrast to DHFR and fibronectin, keratin possesses hierarchical structure ranging from α-helices at the nanometer scale to protofibrils and filaments at the micrometer scale. A turbidity map generated from 24-hour soaking of wool in salt solutions showed that visible signal can only be observed at high concentrations of LiBr (Fig. 1D). FTIR spectra further indicated complete denaturation of both α-keratin from wool and β-keratin from goose feathers, irrespective of their different native configurations (Fig. 1G). Finally, scanning electron microscope (SEM) images of wool before and after treatment with 8 M LiBr, LiCl, and NaBr also corroborate our turbidity measurement findings, where only concentrated LiBr induces denaturation of keratin (Fig. 1J, fig. S4E, and fig. S6).

### Indirect solute effects

Thermodynamically, protein conformational change or denaturation is a complex process where entropy and enthalpy contribute to the overall free energy balance. At lower concentrations of denaturants, entropy and enthalpy balance to favor the native state; however, at higher concentrations, they tend towards the unfolded state (*27*). To elucidate the thermodynamic mechanisms of protein denaturation by LiBr, we conducted isothermal titration calorimetry (ITC) experiments to measure the enthalpic contribution from direct LiBr solute-protein interactions, with commonly used denaturants urea and guanidine hydrochloride (GdnHCl) as a control. Pronounced heat release was detected upon injection of urea or GdnHCl into samples containing DHFR and fibronectin (Fig. 2, A and B, and fig. S7). Enthalpy changes, dependent on protein sizes and exposures of hydrophobic domains (∼-7 kCal mol^-1^ for DHFR and ∼-14 kCal mol^-1^ for fibronectin), correspond to the formation of 1-2 or 3-4 hydrogen bonds per organic denaturant molecule, respectively. These results are consistent with previous studies (*28, 29*), where enthalpy-driven mechanisms of protein denaturation by urea or GdnHCl were established. In contrast, no enthalpy change from direct interaction with proteins was detected when LiBr solution was injected into protein samples under identical protocols, suggesting that solute ions here do not engage in noticeable direct interactions with proteins. Given that LiBr, being a universal and potent denaturant, does not contribute enthalpically to protein free energy in solution, we hypothesize that LiBr shifts the free energy landscape of proteins towards denatured state by affecting its entropy.

**Fig. 2.**
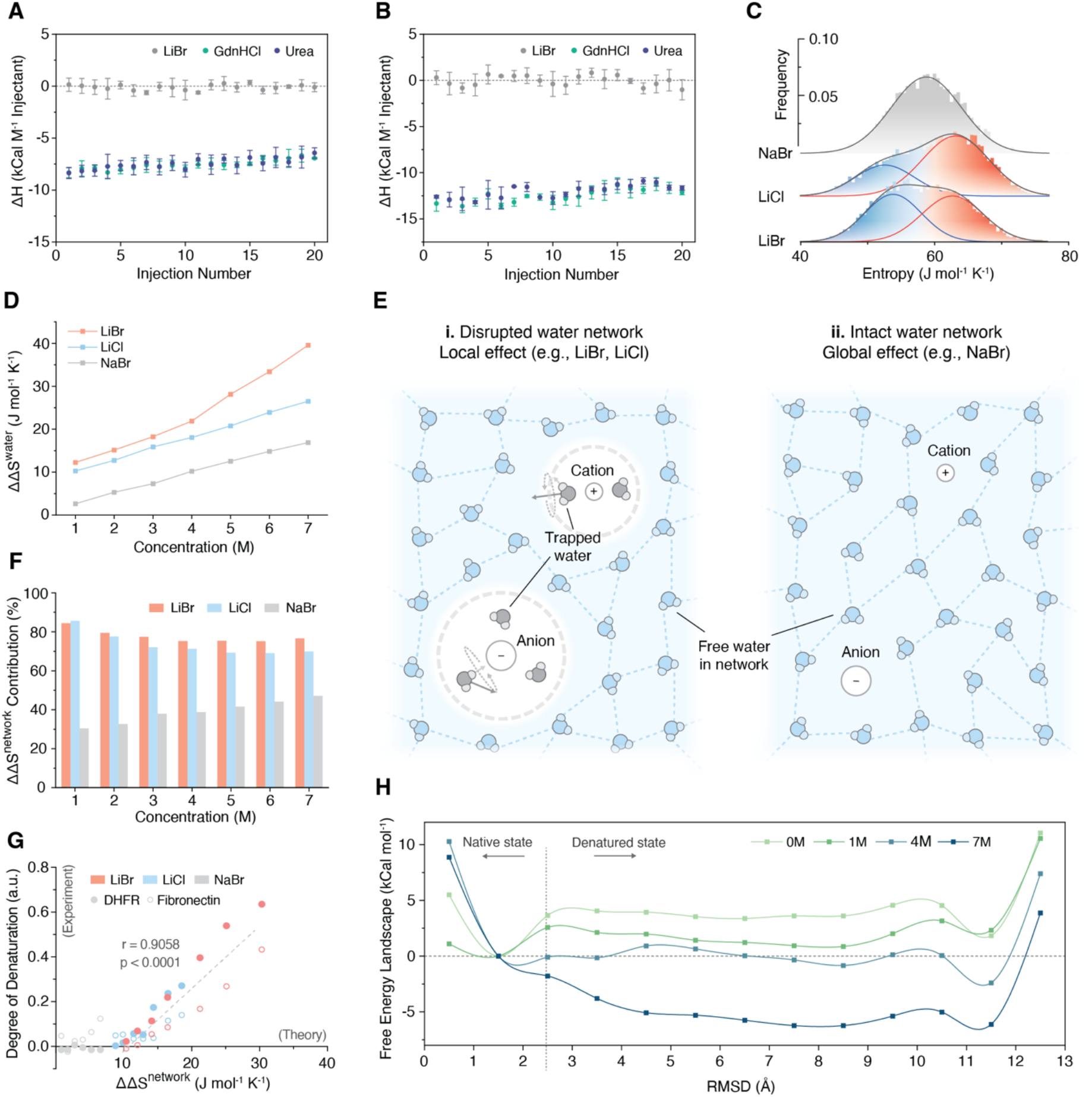
Protein conformational changes come from indirect solute effects manifested as water network entropy. (**A** and **B**), Reaction enthalpy of interactions between proteins (**A**, DHFR, **B**, fibronectin) and denaturants (LiBr, urea, and GdnHCl) as measured by isothermal titration calorimetry. Data are presented as mean ± s.d. (n = 3). (**C**), Distributions of molecular entropy of water molecules in 7 M LiBr, LiCl, and NaBr from MD simulations. (**D**), Total water entropy penalty decrease in different concentrations of LiBr, LiCl, and NaBr solutions. Data are obtained from MD simulations and calculated via the analytical model (Eq. 1). Data are presented as mean ± s.d. (n = 3), however, the error bars are smaller than the data point size. (**E**), Schematics of different ion effects on water structures. For LiBr and LiCl, high charge density of Li^+^ ion results in localized (trapped) water molecules and disrupted water network (**i**), while NaBr induces a more global effect without breaking water network (**ii**). (**F**), Contribution of water network entropy penalty decrease to the total water entropy penalty decrease presented in (**D**). (**G**), Strong correlation between the theoretical value of water network entropy penalty decrease and the protein denaturation ratio observed from FTIR experiments in LiBr and LiCl solutions. Pearson correlation coefficient calculated with a 95% confidence interval for a two-tailed test. Dashed line represents the best fit. (**H**), Free energy landscape of an α-helix peptide (20 amino acids) in pure water and 1, 4, 7 M LiBr from MD simulations.

### Mechanism of entropy-driven protein denaturation

To further test the hypothesis of entropy-driven denaturation by LiBr, molecular dynamics (MD) simulations were employed to provide experimentally inaccessible atomistic-level insights. First, we confirmed that local water density around ions (fig. S8, A and B) in our simulations agrees well with previous X-ray and neutron diffraction experimental results (*30*). The cooperativity of ion hydration (*31*) was also observed, where local water structures are determined by both ions and their counterions. Molecular water entropy, encompassing both translational and rotational entropy of water molecules, was calculated based on TIP3P (*32*) water model using Two-Phase Thermodynamic (2PT) method, which has been validated as the most accurate for replicating experimental entropy values (*33*). In both LiBr and LiCl solutions, two distinct populations of water molecules are detectable (Fig. 2C). As suggested by evidence of local effects of Li^+^ ions (*34*), the subset characterized by lower average entropy is identified as ion-bound water, whereas the other, possessing higher entropy, is identified as free water that constitutes the hydrogen bonded water network. LiBr solution also demonstrates a higher proportion of ion-bound water compared to LiCl solution (Fig. 2E, i, and fig. S8, C and D). This enhanced capacity of Li^+^ ions to sequester water in LiBr could stem from the more effective separation between Li^+^ and Br^-^ ions (*8, 9*), attributable to the lower charge density of Br^-^ ions and the substantial size disparity between Li^+^ and Br^-^ ions. In contrast, the NaBr solution exhibits a single population suggesting to the intact hydrogen bond network of pure water (Fig. 2E, ii, and fig. S8E), which could potentially be explained by the absence of localized high charge density (*9, 31*).

The prominent differences in both water network size and denaturation capability between LiBr, LiCl, NaBr, and pure water, point out to fundamental change in water network entropy as a possible culprit. To further highlight this concept, we established an analytical model describing the decrease of total entropy penalty from water molecules in solution (ΔΔ*S*^water^), which drives the overall free energy balance towards the denatured state.

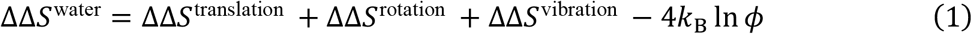

Here, ΔΔ*S*^translation^, ΔΔ*S*^rotation^, and ΔΔ*S*^vibration^ represent the entropy penalty decrease from individual water molecules and -4*k*_B_ ln ϕ denotes the entropy penalty decrease of water as a collective network (ΔΔ*S*^network^), with ϕ representing the ratio of water network sizes in ionic solutions and pure water (see **Supplementary Text** for derivation of eq. 1 and further details).

The total decrease of water entropy penalty from different ion pairs can be calculated using eq. 1 with values of individual contributions obtained from molecular dynamics simulation (Fig. 2D and fig. S9). Notably, the decrease of water network entropy penalty ΔΔ*S*^network^ appears to be the predominant factor over contributions of rotational and translational entropy of individual water molecules, especially for LiBr and LiCl, the two potent protein denaturants (Fig. 2F). This key factor has been overlooked in several previous studies leading to considerable controversy regarding the effects of solutes on water dynamics and protein unfolding (*35, 36*). Furthermore, the theoretically calculated ΔΔ*S*^network^ and the experimentally observed ratio of denatured and native α-helix around 1650-1655 cm^-1^ from FTIR in LiBr and LiCl solutions showed a strong correlation with the Pearson correlation coefficient of 0.9058 (Fig. 2G). In contrast, contributions from rotational and translational entropy exhibited a weaker correlation (fig. S10), with minimal contributions to the total decrease of water entropy and misalignment with the denaturation capabilities of different salt solutions. The strong quantitative agreement between theoretical predictions and experimental results suggests the universal predictive capability of the water network model across various ion types and concentrations, particularly at conditions where water network entropy plays a dominant role. For salts and concentrations where ΔΔ*S*^network^ is less than 10 J mol−^1^ K−^1^, our model predicts no denaturation or indicates that water network entropy is not the primary driving force behind any observed denaturation, as exemplified by NaBr.

Finally, protein free energy landscapes at different LiBr concentrations were constructed using well-tempered metadynamics simulation (*37*) (See **Methods** for details), where a stable α-helix peptide of 20 amino acids was adopted as a model protein (Fig. 2H, and figs. S11 to S15). The increase of ion concentrations shifts the free energy landscapes and lowers the energy barrier for unfolding, which lends further confirmation of the universal denaturation capacity of LiBr. Thus, solute-induced water network disruption potentiates the reduction of entropy penalty when proteins sequester water molecules from bulk water network during unfolding. This facilitates unfolding and potentiates large-scale protein conformational changes and denaturation.

### Keratin regeneration with closed-loop recycling and facile manufacturing

In contrast with conventionally applied enthalpy-driven denaturants (e.g., urea, GdnHCl), the entropy-driven denaturation mechanism can address limitations in protein regeneration, including the waste and pollution from single-use organic chemicals and the complexity of fabrication. Hereby, we developed an alternative protocol for keratin-rich waste regeneration with closed-loop recycling of LiBr solution followed by facile manufacturing based on the entropy-driven denaturation mechanism (Fig. 3A). Protein resources such as animal fragments or wool-based textiles are denatured by an aqueous solution of 8 M LiBr, with heat applied to accelerate reaction kinetics and prevent aggregation during extraction. A small amount of 1,4-dithiothreitol (DTT) is added as a reducing agent to break down the dense disulfide matrix in keratin. After extraction, the mixture is hot filtered, cooled down and separated into a subnatant of denatured keratin gel and a supernatant of LiBr solution driven by spontaneous aggregation (fig. S16, A and B). Since the LiBr does not induce denaturation via direct interactions, the resulting solution can be continuously recycled in a closed loop, which not only eliminates pollution from waste organics but also significantly reduces the consumption and cost of denaturants.

**Fig. 3.**
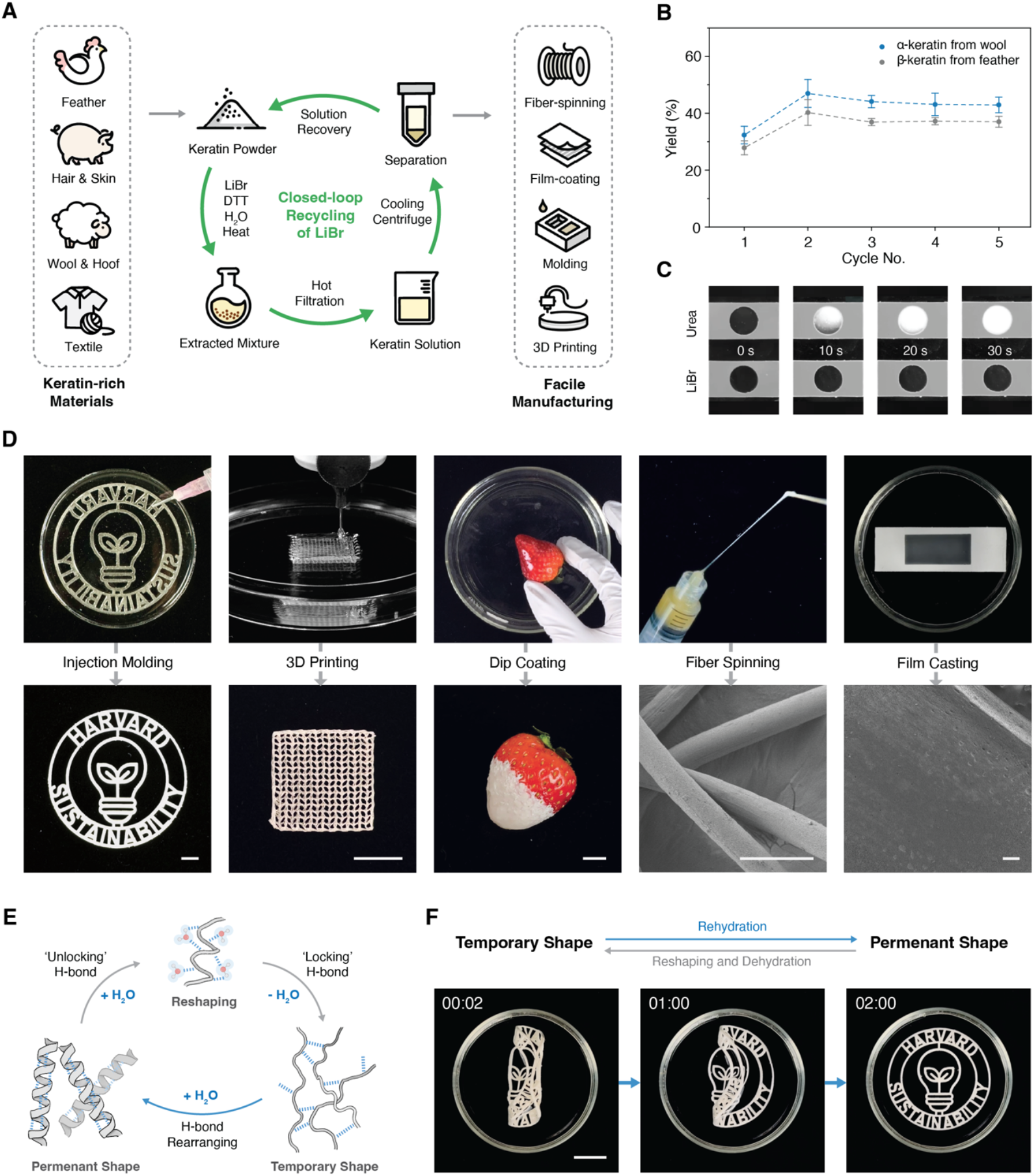
Keratin regeneration with closed-loop recycling and facile manufacturing. (**A**), Schematic showing the regeneration protocol of keratin materials with closed-loop recycling of LiBr solution. (**B**), Extraction yield of α-keratin from wool and β-keratin from goose down. Data are presented as mean ± s.d. (n = 7 for α-keratin, n = 3 for β-keratin). (**C**), Immersing extracted keratin in transparent well spacers in water. Keratin gel from LiBr extraction quickly turns into a white solid phase, while keratin solution extracted with urea remains soluble. (**D**), Facile manufacturing strategies including injection molding, 3D printing, dip coating, fiber spinning, and membrane casting. Scale bars, 1 cm for optical images, 200 μm for SEM images. (**E**), Schematic of the shape memory effect triggered by hydration signal. (**F**), Demonstration of shape memory effect by placing a dehydrated keratin sample in water. Scale bar: 3 cm.

To test the cyclic stability of the closed-loop recycling protocol, we performed multiple iterations of extraction using both α-keratin from wool and β-keratin from goose feathers. The yield remains consistently around 40% after five cycles of extraction (Fig. 3B, and fig. S17), while the first cycle exhibits a slightly lower yield possibly due to a small amount of keratin remaining in the solution after separation. Unchanged composition of the recycled LiBr solution is further confirmed with thermogravimetric analysis (TGA) and coupled Fourier transform infrared spectroscopy (TG-FTIR) of the emerging gas (fig. S18).

Notably, in the absence of direct binding to denaturants, the spontaneous aggregation of denatured keratin enables a simple separation of condensed gel from the solution, alleviating difficulties with conventional organic reagents (fig. S16C), which usually involves dialysis and lyophilization. Additionally, the separated keratin can undergo a fast phase transition from denatured gel state to renatured solid state when transferring back into water (Fig. 3C, and movie S1), while keratin extracted with organic denaturants remains non-processable until fully dialyzed and lyophilized. The viscous property of the gel and the fast phase transition process enable a wide array of manufacturing modalities including injection molding, membrane casting, dip coating, fiber spinning, and 3D printing (Fig. 3D, and fig. S19) from a broad range of keratin sources (fig. S20) without any additional usage of organic solvents.

### Mechanical tunability and shape-memory of regenerated keratin

Keratins, particularly α-keratins, contain multiple cysteine residues, which form a dense network of intermolecular disulfide bonds. After extraction, these disulfide bonds are reduced to thiol groups, thereby endowing the regenerated keratin material with adjustable covalent crosslinking through oxidation. For instance, low level of crosslinking allows the α-helices to rearrange more freely under stress. As a result, reduced keratin preserves a high flexibility with a modulus of 124.3 ± 18.8 kPa and a fracture strain of 435 ± 61% (fig. S21, A to C). On the contrary, formation of multiple disulfide bonds can significantly increase the elastic modulus of oxidized keratin by an order of magnitude to 1.35 ± 0.11 MPa and decrease the fracture strain to 95 ± 9% (fig. S21, D to F). Raman spectroscopy of keratin samples under 50% strain suggests that most secondary structures remain unchanged within reduced keratin upon stretching, whereas a more discernible loss of α-helix domains and an increase of β/random structures can be observed in oxidized keratin (fig. S22). Differences between the two conditions reveal the effect of disulfide crosslinking on the α-helix network of regenerated keratin, providing the material with tunable mechanical properties from ductile to elastic (fig. S21, G to I).

Additionally, the capability of forming a homogenous object through the fast phase transition in water and the preservation of the native α-helix structure provide a hydration-responsive shape-memory property for the regenerated keratin. Prior studies of Miserez et al. (*38*) and Cera et al. (*15*) revealed that uniaxial strain can induce reversible uncoiling of α-helix into metastable β-like structures within anisotropic bio-based elastomer. Similarly, in the disulfide crosslinked keratin material, the transition between coiled helix and uncoiled strands can also lead to a shape-memory effect facilitated by formation and decay of hydrogen bonds (*39*). Dynamic reformation of hydrogen bonds in a hydrated environment enables the reshaping of the object into a temporary shape without breaking, which can then be locked temporarily through dehydration due to the formation of intermolecular hydrogen bonds. Upon rehydration, the presence of free water molecules can unlock the hydrogen bonds again and facilitate the recovery of native α-helix structures and the permanent shape (Fig. 3E). Notably, this property has been observed exclusively in natural keratin or keratin materials obtained using the LiBr regeneration strategy. In contrast, traditional extraction methods generally result in dispersed powders or compromised secondary structures (fig. S23, and table S1). The shape-memory capability of regenerated keratin material is demonstrated both as a standalone object and as components of complex tensegrity structures (Fig. 3F, figs. S24 and S25, and movies S2-4).

## Discussion

The effect of water networks on protein conformation and stability has long been a subject of extensive debate. Building on the foundational work dating back to the 19^th^ century ‘Hofmeister Series’ and continuing to the present, we investigated the molecular mechanisms underlying hierarchical protein conformational changes in the presence of three monovalent salts—LiBr, LiCl, and NaBr. Notably, our ITC experiments revealed no direct interactions between proteins and ionic solutes despite the significant potency of LiBr as a universal denaturant. This result provides direct evidence of a new mechanism of denaturation that stands in contrast to the established mechanism of denaturation driven by direct binding of organic denaturants to proteins. Instead, it points to an alternative pathway driven by entropy rather than enthalpy.

Thermodynamic and spectroscopic analyses, in combination with atomistic molecular simulations, have shown that large-scale protein conformational changes and denaturation by LiBr and other ionic solutes are mediated by reduction of entropy penalty (Fig. 4). Localized strong solute-water interactions for LiBr, and, to a lesser extent, LiCl, disrupt the water network and reduce the entropy penalty that proteins incur when sequestering water molecules from bulk water phase during denaturation. In contrast, the integrity of water network is largely preserved in NaBr solution, thereby consistently retaining protein native structures across varying concentrations. This entropic mechanism of protein denaturation by salts like LiBr opens new opportunities in technological development of protein extraction from natural sources by alleviating the need in complex and environmentally hazardous steps of separation of organic denaturants from bulk proteins.

**Fig. 4.**
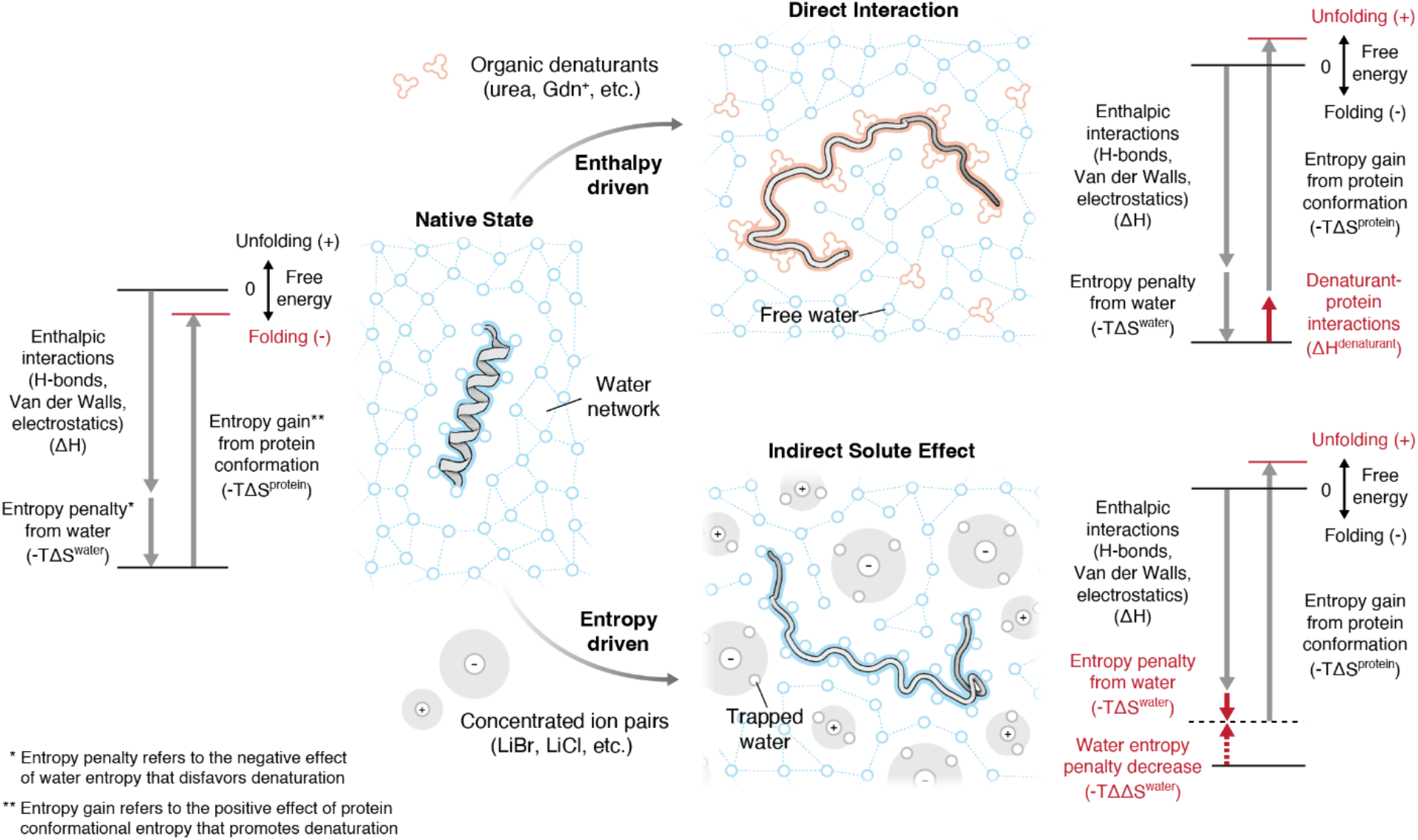
Mechanistic comparison of enthalpy-driven denaturation and entropy-driven denaturation. In native state, proteins adopt a folded conformation driven by the combined effect of enthalpy and entropy. Traditional organics denaturants, such as urea and Gdn^+^, interact directly with proteins, introducing additional enthalpic contributions that shift the equilibrium toward the unfolded state. In contrast, concentrated ion pairs, such as LiBr, disrupt the water network, decreasing the entropy penalty associated with protein-water interactions, and thereby achieve protein denaturation indirectly through an entropy-driven mechanism.

## Data availability

All the source data can be readily accessed via https://doi.org/10.7910/DVN/U6HAQS

## Code availability

All the customized codes used for data analysis are available at https://github.com/junlang-liu/Entropy_code

## Acknowledgements

We thank M. Rosnach for photography and illustrations, S. Mondal for suggestions on molecular simulations, K. Park, E. Serebryany and A. Bitran for discussions on water structure and protein denaturation mechanisms, H. Chang and J. F. Zimmerman for discussions on material properties. K. Shani and Y. Jang for discussions on biological properties and tissue engineering potential of keratin. This work was sponsored by National Institutes of Health (R35GM139571 and R01EY030444 to E.I.S), the John A. Paulson School of Engineering and Applied Sciences at Harvard University (K.K.P.), the National Science Foundation through the Harvard University Materials Research Science and Engineering Center (DMR-2011754 to K.K.P.), the Health@InnoHK program of the Innovation and Technology Commission of the Hong Kong SAR Government, and the Medical & Health Informatics Laboratories at NTT Research, Inc. This work was performed in part at the Harvard University Center for Nanoscale Systems (CNS), a member of the National Nanotechnology Coordinated Infrastructure Network (NNCI), which is supported by the National Science Foundation under NSF award no. ECCS-2025158. The content is solely the responsibility of the authors and does not necessarily represent the official views of NTT Research, Inc.

## Author contributions

Y.W., J.L., K.K.P., and E.I.S. designed and developed the project. Y.W. and J.L. performed turbidity measurements, tryptophan fluorescence, and FTIR. Y.W., J.L., and D.W. performed ITC experiments. Y.W., J.L., and S.C. performed Raman experiments. J.L. did *E. coli* cell culture, DHFR protein purification and quality control. J.L. conceived the entropy-driven model and developed the analytical model. J.L. conducted all the simulations and related data analysis. Y.W. performed TG-FTIR, polarized Raman, DLS, and TGA measurements. Y.W. performed keratin extraction, mechanical tests, and demonstrations. Y.W. and M.M.P. fabricated keratin fibers and performed SEM imaging. Y.W. and R.I. did SDS-PAGE. Y.W. and J.L. prepared the initial manuscript. Y.W., J.L., K.K.P., and E.I.S. revised the manuscript, with contributions from all the other authors. K.K.P. and E.I.S. supervised the research.

## Supplementary materials

Materials and Methods

Supplementary Text

Figs. S1 to S25

Tables S1 and S2

**Correspondence and requests for materials** should be addressed to Eugene I. Shakhnovich or Kevin Kit Parker

