## Supplementary Materials for "Entropy Changes in Water Networks Promote Protein Denaturation"

### Materials and Methods

Escherichia coli dihydrofolate reductase expression

Cell culture. *E. coli* dihydrofolate reductase (DHFR, UniProt: P0ABQ4) (wild-type or W30C mutant) was overexpressed in BL21(DE3) *E. coli* cells. The plasmid was synthesized by GenScript Biotech with pET-28a(+) vector and C-terminus 6xHis-tag. All the cell culture were conducted in Terrific Broth (TB) medium (BD Difco™), supplemented with 0.4% (v/v) glycerol (RPI), 4 g $L^{-1}$ glucose (Sigma-Aldrich), 25 mM MOPS (Sigma-Aldrich, pH=7.2), 0.4 mM citric acid (VWR Chemicals BDH®), 40 µl $L^{-1}$ 1% (w/v) ferric ammonium citrate (AMRESCO) solution, and 50 µg $ml^{-1}$ kanamycin (Millipore-Sigma). All the concentrations denoted here are final concentrations. 2% (v/v) frozen cell stock solution of BL21(DE3) *E. coli* cells transformed with DHFR (wild-type or W30C mutant) plasmid was used for a small overnight (~12 h) culture at 37 °C with shaking at 250 rpm. 2% (v/v) overnight culture solution was used to start the production culture, initially at 37 °C with shaking at 250 rpm. After 3.5 hours at which the cell growth started the mid-log phase with an optical density of around 0.6 at 600 nm, isopropyl β-D-thiogalactopyranoside (IPTG, RPI) was added to a final concentration of 100 µM to induce the protein overexpression. The temperature was decreased to 18 °C, with shaker speed kept as 250 rpm. The cell pellets were harvested after 12-hour low-temperature protein overexpression through centrifugation.

Protein purification. Three different buffers were used in DHFR protein purification: Lysis buffer [100 mM bicine (Sigma-Aldrich), 5 mM imidazole (Millipore-Sigma OmniPur®), 250 mM NaCl (VWR Chemicals BDH®), pH=9]; Buffer A [20 mM Tris (J.T. Baker™), 5 mM imidazole, 250 mM NaCl, pH=8.5]; Buffer B [20 mM Tris, 50 mM imidazole, 250 mM NaCl, pH=8]. Every 1 g cell pellet (from 50 ml cell culture solution) was resuspended in 1.5 ml lysis buffer, supplemented with 0.01% (v/v) Nuclease (Millipore-Sigma Benzonase®), 1% (v/v) Protease Inhibitor Cocktail Set II (Millipore-Sigma Calbiochem®), and 1 mg $ml^{-1}$ lysozyme (Thermo Fisher Scientific). All the concentrations denoted here are final concentrations. The resuspended solution was then sonicated, centrifuged with supernatant collected. cOmplete™ His-Tag Purification Resin (Roche) was used to further isolate DHFR protein from supernatant in gravity flow columns (Bio-Rad). Buffer A was first used to wash away non-bound proteins and other cell debris, followed by elution of Buffer B, where DHFR was washed out and collected. Only the fractions with $A_{280}/A_{260} > 1.7$ were pooled based on the absorbance measurement with a NanoDrop spectrophotometer (Thermo Fisher Scientific). DHFR was then washed into 20 mM Tris buffered saline (pH=7.0, with 250 mM NaCl) and concentrated into a final concentration of 3 mg $ml^{-1}$ using ultrafilter (Sartorius Vivaspin®). Mass spectrometry (Bruker Impact II q-TOF) was used to confirm that DHFR (both wild-type and W30C mutant) was successfully synthesized and purified, after which protein solution was aliquoted and stored in -70 °C.

Turbidity assay

Turbidity measurements at $OD_{405}$ were performed using a Biotek PowerWave HT 340 microplate reader. For each condition, protein solutions was mixed with solutions of LiBr, LiCl, and NaBr (Sigma-Aldrich) to a final concentration of 0.4 mg $ml^{-1}$ for DHFR or 0.1 mg $ml^{-1}$ for human fibronectin (BD Biosciences), with a total volume of 100 µl in a 96-well plate. Samples were then immediately placed into the microplate reader and $OD_{405}$ was continuously measured for 1 h at room temperature. For keratin, 80 mg of cleaned wool (R. H. Lindsay Wool Company) was soaked into 4 ml of salt solutions with 0.1 M DTT (Sigma-Aldrich) in quartz cuvettes, then heated to 70 °C to accelerate the denaturation kinetics. Samples of the solutions were taken at 24 h and 48 h, added into 96-well plate, and cooled at room temperature before $OD_{405}$ measurement. Photos recording the condition of the solution were taken after cooling for 1h.

FTIR and TG-FTIR

All FTIR measurements were conducted using a Nicolet iS50 FTIR spectrometer. Protein solutions with final concentrations of 1 mg $ml^{-1}$ DHFR, 0.5 mg $ml^{-1}$ fibronectin, and 1 mg $ml^{-1}$ keratin (α-keratin from Angora wool, R. H. Lindsay Wool Company; β-keratin from goose down, Dream Solutions USA), respectively, were prepared with a gradient of LiBr concentrations ranging from 0 to 8 M, then stabilized for at least 3 h prior to measurements. During acquisition, background information was initially recorded by applying pure LiBr solutions onto the ATR crystal, which was then replaced by the corresponding protein solutions, with a total of 64 scans collected for each sample. Solid samples, including regenerated keratins and raw materials, were measured using the same ATR-FTIR setup. Gas phase FTIR coupled with TGA was collected using the same equipment connected to the Discovery TGA 550. Data analysis and background subtraction were carried out with OMNIC v.9.2.86 software. To calculate the ratio of protein denaturation for the correlation analysis in Fig. 2G and Fig. S10, we measured the intensity change at 1652.5 $cm^{-1}$, corresponding to the denaturation of α-helix structure.

Raman Spectroscopy

Raman experiments were performed with a Horiba LabRam HR Evolution Raman confocal system (633nm excitation, 50x, 0.5NA, 13mW). For DHFR experiment, a customized imaging chamber made by sandwiching a 1 mm thick PDMS (Sylgard™ 184, Dow Corning) well between two glass slides (fig. S2) was loaded with a solution of 3 mg $ml^{-1}$ DHFR and 2 M LiBr. Each spectrum was collected with 600 gr $mm^{-1}$, 180 s acquisition time and 2 accumulations. For solid samples, 1800 gr $mm^{-1}$, 60 s acquisition and 4 accumulations were used. Data collection and analysis were carried out with Labspec v.6.5.1 software.

Dynamic light scattering

DLS experiments were performed using a Malvern Zeitasizer Pro system. Fibronectin in LiBr solutions were prepared with a final concentration of 0.5 mg $ml^{-1}$, stabilized for at least 3 h, and filtered through a 0.22 µm PTFE filter (VWR) before transferred into 40 µl cuvettes (Malvern Panalytical) for measurement. Keratin solutions after extraction were diluted to approximately 1 mg $ml^{-1}$, filtered, then transferred into 1 ml cuvettes (Malvern Panalytical). Samples were heated to 70 °C for 30 min and stabilized at designated temperature for 10 min before measurement. Data collection and analysis were carried out with ZS Xplorer v.3.0.0.53 software.

Scanning electron microscopy

Samples were mounted on a 12.5 mm diameter SEM stub covered with carbon tape, then sputtered-coated with Pt/Pd with an EMS 150T ES sputter coater with 10 nm thickness. SEM images were taken with a Zeiss Gemini 360 field emission scanning electron microscope with an electric high tension of 3 kV and SE2 detector.

Isothermal titration calorimetry

ITC experiments were conducted using a Malvern Panalytical MicroCal VP-ITC. During the experiments, the sample chamber was filled with 1.8 ml of 1 mg $ml^{-1}$ DHFR (W30C mutant) solution or 0.5 mg $ml^{-1}$ fibronectin solution, while the syringe was loaded with 100 μM solution of LiBr, urea, or GdnHCl (Sigma-Aldrich). Corresponding control experiments were performed by titrating milli-Q water into the same concentration of protein solution to measure and subtract the dilution heat of proteins. Additionally, parallel control groups were also performed by titrating 100 μM ligands into a chamber filled with milli-Q water to measure the dilution heat of ligands, which were found to be significantly smaller than the detectable binding enthalpy between proteins and ligands (fig. S7). Each individual run comprised titrating 7 μl ligand solution 25 times into the chamber, with a duration of 12 s, and interval of 240 s, a filter period of 2 s, a stir speed of 309, and temperature controlled at 30 °C. Data collection and analysis including baseline adjustment and integration of enthalpy were carried out with VPViewer2000 v.1.29.1 software.

Molecular dynamics simulations

General simulation setup. All the molecular dynamics simulations were conducted with NAMD 2.14 (*40*) and analyzed and visualized with VMD 1.9.3 (*41*), unless otherwise denoted. Amber ff14SB force field (*42*) were used for atomistic protein simulation, combined with TIP3P water model (*43*) for explicit solvation and Li/Merz ion parameters (*44*) for monovalent ions. All initial configurations of simulation box were prepared using tLEaP program (*45*), followed by minimization, heating, constant pressure (NPT) equilibration, and constant volume (NVT) equilibration. All the production simulations were run under constant particle number, constant volume, and constant temperature (i.e., canonical ensemble (NVT)). Periodic boundary condition was applied to prevent the boundary effect. Langevin thermostat with damping coefficient of 1 $ps^{-1}$ was used to control simulation temperature as 298 K, without coupling hydrogen atoms. Electrostatic and van der Waals interactions were cut off beyond 12 Å. Particle Mesh Ewald (PME) method was employed to deal with long-range electrostatic interactions. Rigid bond constraints between hydrogen and any other atoms were applied through SHAKE/RATTLE algorithm. The integration time step was set as 2 fs with short-range nonbonded forces updated every 2 fs and long-range electrostatic forces updated every 4fs.

Initial configuration preparation and equilibration. To ensure that total number of ions and water molecules are correctly determined for all salts and concentrations, we first decided the simulation box sizes and thus ion numbers. All the simulation boxes were fixed to the size with three dimensions around 50 Å. Water molecules were initially placed using tLEaP program (*45*), the numbers of which were then adjusted according to volume changes after 100 ps minimization, 200 ps heating, and 2 ns constant pressure (NPT) equilibration. After the water molecule number corrections, another round of 100 ps minimization, 200 ps heating, 2 ns constant pressure (NPT) equilibration, plus 2 ns constant volume (NVT) equilibration, were conducted to ensure the correct setup and complete equilibration of simulation systems.

Water structure characterization. The equilibrated initial configurations containing water molecules and ions were prepared as stated above. The production simulations for water structure characterization were run for 100 ns with trajectories recorded every 10 ps. The radial distribution function $g(r)$ between cations (or anions) and oxygen atoms of water can be used to characterize local water structure around ions. It was calculated via VMD and averaged among all cations (or anions) and all trajectory frames with the definition as

$$g(r) = \frac{dn_r}{4\pi r^2 \cdot dr \cdot \rho_{\text{bulk}}} \quad \text{(Eq. S1)}$$

where $dn_r$ is the number of oxygen atoms of water within the shell of thickness at the distance $r$, $4\pi r^2 \cdot dr$ is the volume of the shell, $\rho_{\text{bulk}}$ is the bulk average density of oxygen atoms of water in the simulation box.

Water entropy calculation. The equilibrated initial configurations containing water molecules and ions were prepared as stated above. The production simulations for water entropy calculation were run for 20 ps (*33*) with trajectories recorded every 2 fs. All the water entropy calculations were conducted based on Two-Phase Thermodynamic (2PT) model (*33*, *46*) using corresponding open source program (*47*). The resulting histograms were fit with two Gaussian distributions with no boundary conditions. Initial values of peak centers were determined using local minima of 2$^{nd}$ derivative. For LiBr and LiCl, fitting achieved convergence with Chi-Square tolerance value of $10^{-6}$. For NaBr, one peak will collapse to zero and only a single peak remained upon convergence.

Protein free energy landscape construction. To examine changes in free energy that drive denaturation, we used metadynamics simulation to explore the free energy landscape of the complex molecular system including water, ions, and protein. A short alanine-based peptide with 20 amino acids (AAAAKAAAAKAAAAKAAAAK), which has been confirmed experimentally (*48*) to have a stable alpha helix structure in pure water, was adopted as the model protein to study the difference of protein free energy landscapes under different concentrations of LiBr. We used ColabFold v1.5.5 (*49*) to obtain the initial atomistic protein structure, with N-terminus acetylated and C-terminus amidated using PyMOL 2.5.5. The equilibrated initial configurations containing protein, water molecules and ions were prepared as stated above. Well-tempered metadynamics simulations were implemented for protein free energy landscape construction using Colvars module (*37*) on top of standard molecular dynamics simulations as stated above. Root mean square displacement (RMSD) from the backbone of native alpha helix protein structure was selected as the collective variable to guide the sampling of unfolded confirmations. After the first run of metadynamics simulation, two more protein conformations (fig. S11A) with medium RMSD (6 Å) and large RMSD (12 Å) respectively, were chosen as two other start structures. In total, nine metadynamics simulations for each LiBr concentration, started from three different initial protein conformations, were conducted. Each simulation was run for around 2 μs with trajectory recorded every 100 ps (fig. S9, B and C). Convergence of metadynamics simulations was examined using block analysis via the open-source, community-developed PLUMED library, version 2.8.3 (*50*).

Rigorous criteria have been implemented to ensure the accuracy of free energy landscapes derived from metadynamics simulations (fig. S12-S15). Initially, simulations are conducted until the collective variable (CV) iterates over its entire possible range, in this case, 0 to 12.5 Å (first column). Secondly, the final height of the Gaussian bias potential added at each CV position must be less than $5*10^{-3}$ kCal $M^{-1}$ (second column). Lastly, the convergence of the simulations is further confirmed by observing a plateau in the average standard deviation of the potential of mean force (PMF) as calculated from block analysis, with increasing block sizes (third column). Six converged simulations out of nine for each LiBr concentration were used for final protein free energy landscape construction.

Keratin extraction protocol

The pretreatment of keratin-rich sources (Angora wool, Merino wool shirt, and feathers from the R. H. Lindsay Wool Company, Merino Tech, and Dream Solutions USA, respectively. Hair samples collected from local barber shop.) involves washing with ethanol (VWR 200 proof) in a Soxhlet extractor for 48 h to remove organic residues, followed by rinsing with water and allowing the material to dry at room temperature overnight. Subsequently, long wool fibers were cut into smaller fragments (less than 1 cm in length) to facilitate dispersion in solution. During the extraction process, 10 g of keratin fragments were suspended in a 150 ml aqueous solution containing 8 M LiBr and 0.1 M DTT. The suspension was stirred for 36 h in an insulated environment at 90°C. Afterwards, insoluble residues were filtered through cotton cloth with 80/180 μm mesh size while still hot, then stored at 4°C overnight. A viscous, gel phase of aggregated keratin was observed, and centrifugation (3,000 r.p.m., 4°C) further separated the mixture into an upper phase of LiBr solution and a lower phase of keratin gel. The LiBr solution could then be easily collected and reused in the following extraction cycle along with a new batch of keratin source and DTT. A small amount of 8 M LiBr stock solution were added to compensate the solution loss from filtration process in order to maintain the total volume at 150 ml.

The keratin content of the aggregated gel was measured by taking 1 ml of gel and allowing it to fully solidify in water, freeze-dried, then lyophilized to obtain a dehydrated solid and measure its weight. It was found that 1 ml of keratin gel contains 392.8 ± 18.6 mg of α-keratin or 347.9 ± 23.7 mg of β-keratin (n = 6). To calculate the extraction yield, the volume of the keratin gel obtained from each batch was measured, multiplied by the corresponding content, and divided by the initial weight of keratin fragments. The denatured keratin gel could be stably stored at 4°C in the absence of oxygen, with a shelf life longer than 6 months in absence of oxygen.

Thermogravimetric analysis

TGA of the LiBr solution prior to the 1$^{st}$ cycle and the recycled LiBr solution after the 5$^{th}$ cycle were conducted with a Discovery TGA 550. Around 30 mg of solution was loaded onto a platinum HT sample pan, stabilized under nitrogen flow for 10 min before measurement. Experiments were performed by heating samples to 100 °C at a rate of 5 °C min$^{-1}$, then to 600 °C at a rate of 10 °C min$^{-1}$ in an air flow of 90 ml min$^{-1}$. The mass of the sample was continuously measured while the evolving gas from the sample was analyzed simultaneously by TG-FTIR. Data collection and analysis were carried out with TRIOS v.5.2.2 software.

Keratin sample preparation

All samples were regenerated using α-keratin gel unless otherwise noted. The reverse mold for injection molding was created by casting PDMS (Sylgard™ 184, Dow Corning) around a 3D printed logo. Keratin gel was injected into the mold and transferred into a water bath until solidified. Membrane casting and dip coating followed a similar process by casting a layer of gel on glass coverslip or coating targets, then transferred into a water bath. Fiber spinning was achieved by injecting the gel through a 21 gauge needle (BD PrecisionGlide™) at a constant rate of 0.1 ml min$^{-1}$ into a bath of 0.4 M $NaH_2PO_4$ (Sigma-Aldrich) solution to accelerate the phase transition process. 3D printing was carried out using a Cellink BIO X 3D printer. The keratin gel was extruded into a supporting bath of 25% w/v Pluronic F127 (BASF) under an absolute pressure of 50 kPa and through a 22 gauge needle (Nordson EFD Precision Tips) that was moving at a speed of 4 mm s$^{-1}$. The Pluronic bath was then removed by cooling and rinsing with water.

Keratin samples for mechanical tests, Raman and polarized Raman spectra were prepared by film casting a layer of keratin gel onto glass coverslips with a thickness around 0.4 mm, then transferred into a water bath to solidify the gel. Oxidized samples were fabricated by further transferring into a solution of 1% $H_2O_2$ for 1 h to reconnect the disulfide bonds. Finally, samples were sectioned into 5 × 20 mm rectangles and stored in water prior to measurements.

The tensegrity structure for the shape memory demonstration was created by oxidizing keratin films in a curled permanent state to as compression units. These units were then interconnected with nylon threads (Singer®) to serve as tension units (fig. S21). Samples were fixed in a distorted temporary state by desiccating for 90 min with addition of external stress.

Mechanical tests

Mechanical tests were performed with a CellScale biaxial tester with 2.5 N load cells. For cyclic stretching curves, samples were tested at a strain rate of 5% per second to a maximum of 50% for 10 cycles. For yield strain measurements, samples were tested at a strain rate of 5% until fracture. Sample thickness was measured with a Mitutoyo Absolute Digimatic digital caliper. The modulus of reduced and oxidized samples was calculated within the 0-50% strain region.

SDS-Page analysis

SDS-Page analysis of the extracted keratin was performed with 4–15% Precast Gels (Mini-PROTEAN® TGX™) placed in Mini Trans-Blot® Cell connected to PowerPac™ Basic Power Supply for 40 minutes at 200V. We followed the instructions provided by Bio-Rad and used SDS buffer (10xTris/Glycine Buffer), 4x Laemmli sample buffer, and Precision Plus Protein™ Kaleidoscope™ Prestained Protein Standards as a ladder buffer. All the devices and the reagents were purchased from Bio-Rad. Type I keratin with a molecular weight around 60 kDa and type II keratin with a molecular weight around 40 kDa can be observed (fig. S15).

Tryptophan fluorescence measurements

Tryptophan (Trp) fluorescence measurements of DHFR in LiCl were performed using a Cary Eclipse fluorescence spectrophotometer in a 1-cm path-length quartz cuvette. Protein solutions with a final concentration of 0.1 mg $ml^{-1}$ DHFR were prepared with 20 mM Tris buffered saline (pH 7.0, with 250 mM NaCl) and a gradient of LiCl concentrations ranging from 0 M to 8 M. The fraction of denatured protein was calculated by normalizing the ratio of intensity at 340 nm (Trp buried in a hydrophobic core, indicating that DHFR remains folded) and 357 nm (Trp quenched by solvent, indicating denaturation). Data collection and analysis were carried out with Cary Eclipse Scan Application v.1.1. software.

**Supplementary Text**

Analytical model of entropy penalty decrease

The protein unfolding reaction is defined as

$$F \leftrightarrows U \qquad \text{(Eq. S2)}$$

where $F$ is the folded state and $U$ is the unfolded state. For folded proteins, since they can maintain stable structures in pure water solution, the free energy change $\Delta G_{\text{pure}}$ of the unfolding reaction in deionized pure water should be positive, i.e.,

$$\Delta G_{\text{pure}} > 0 \qquad \text{(Eq. S3)}$$

According to the unfolding experiments presented in this work, LiBr has the strongest capability of unfolding (denaturing) proteins, followed by LiCl. NaBr cannot denature those tested proteins in this work. Therefore, the free energy changes of the unfolding reaction in different solution environments can be ranked as follow,

$$\Delta G_{\text{LiBr}} < \Delta G_{\text{LiCl}} < \Delta G_{\text{NaBr}} < \Delta G_{\text{pure}} \qquad \text{(Eq. S4)}$$

where $\Delta G_{\text{LiBr}}$ and $\Delta G_{\text{LiCl}}$ for some proteins in high concentration salt solutions are negative (i.e., the unfolding reaction is favored).

The difference between free energy changes of the unfolding reaction in salt solutions $\Delta G_{\text{salt}}$ and that in pure water $\Delta G_{\text{pure}}$ can be written as

$$\begin{aligned}\Delta\Delta G &= \Delta G_{\text{salt}} - \Delta G_{\text{pure}} < 0 \\ &= \Delta H_{\text{salt}} - \Delta H_{\text{pure}} - T\Delta S_{\text{salt}} - \left(-T\Delta S_{\text{pure}}\right) \\ &= \Delta\Delta H - T\Delta\Delta S\end{aligned} \qquad \text{(Eq. S5)}$$

where enthalpy change difference $\Delta\Delta H$ equals to 0 given the ITC data (Fig. 3, A and B) that no direct interaction between proteins and ions was observed. Thus, we have

$$\Delta\Delta G = -T\Delta\Delta S < 0 \qquad \text{(Eq. S6)}$$

where $\Delta\Delta S$ is the entropy penalty decrease due to high concentration salts and should be positive.

Entropy penalty decrease caused by high concentration salts can be written as

$$\begin{aligned}\Delta\Delta S &= \Delta S_{\text{salt}} - \Delta S_{\text{pure}} > 0 \\ &= \Delta S_{\text{salt}}^{\text{protein}} + \Delta S_{\text{salt}}^{\text{water}} - \left(\Delta S_{\text{pure}}^{\text{protein}} + \Delta S_{\text{pure}}^{\text{water}}\right) \\ &= \Delta S_{\text{salt}}^{\text{water}} - \Delta S_{\text{pure}}^{\text{water}}\end{aligned} \qquad \text{(Eq. S7)}$$

where protein conformational entropy changes $\Delta S^{\text{protein}}$ during unfolding is independent of solutions and thus cancel with each other, $\Delta S^{\text{water}}$ is the entropy change of water in the solution.

The entropy change of water molecules in the solution $S^{\text{water}}$ is composed of two parts, i.e., molecular entropy and network entropy. Molecular entropy is composed of translational, rotational, and vibrational entropy of individual water molecules themselves. Network entropy comes from the collective behavior of water network, which considers different combinations water molecules can have to form a network. Therefore, the total entropy change of water during protein unfolding can be expressed as

$$\Delta S^{\text{water}} = \Delta S^{\text{translation}} + \Delta S^{\text{rotation}} + \Delta S^{\text{vibration}} + \Delta S^{\text{network}} \quad \text{(Eq. S8)}$$

Consider a water network with $N$ water molecules (so-called *free* water molecules since they are not bound by either proteins or ions), the number of microstates for each water molecule inside the network can be expressed as

$$\Omega = \left(N^{\text{free}} - 1\right)\left(N^{\text{free}} - 2\right)\left(N^{\text{free}} - 3\right)\left(N^{\text{free}} - 4\right) \quad \text{(Eq. S9)}$$

given one water molecule can form four hydrogen bonds with other water molecules in the network.

Then the network entropy of each water molecule is

$$S^{\text{network}} = k_{\text{B}} \ln(\Omega) = k_{\text{B}} \ln\left(\left(N^{\text{free}} - 1\right)\left(N^{\text{free}} - 2\right)\left(N^{\text{free}} - 3\right)\left(N^{\text{free}} - 4\right)\right) \quad \text{(Eq. S10)}$$

where

$$\Delta S^{\text{network}} = -k_{\text{B}} \ln\left(\left(N^{\text{free}} - 1\right)\left(N^{\text{free}} - 2\right)\left(N^{\text{free}} - 3\right)\left(N^{\text{free}} - 4\right)\right) \quad \text{(Eq. S11)}$$

The entropy penalty decrease originated from the shrinkage of intact water network size in high salt concentration systems can be expressed as

$$\begin{aligned}
\Delta\Delta S^{\text{network}} &= \Delta S^{\text{network}}_{\text{salt}} - \Delta S^{\text{network}}_{\text{pure}} \\
&= -k_{\text{B}} \ln\left(\frac{\left(N^{\text{free}}_{\text{salt}} - 1\right)\left(N^{\text{free}}_{\text{salt}} - 2\right)\left(N^{\text{free}}_{\text{salt}} - 3\right)\left(N^{\text{free}}_{\text{salt}} - 4\right)}{\left(N^{\text{total}}_{\text{pure}} - 1\right)\left(N^{\text{total}}_{\text{pure}} - 2\right)\left(N^{\text{total}}_{\text{pure}} - 3\right)\left(N^{\text{total}}_{\text{pure}} - 4\right)}\right) \\
&\doteq -4k_{\text{B}} \ln\left(\frac{N^{\text{free}}_{\text{salt}}}{N^{\text{total}}_{\text{pure}}}\right) \\
&= -4k_{\text{B}} \ln \phi
\end{aligned} \quad \text{(Eq. S12)}$$

where $N^{\text{total}}_{\text{pure}}$ is the total number of water molecules in pure water systems as all water molecules here are considered as *free* , both $N^{\text{free}}_{\text{salt}}$ and $N^{\text{total}}_{\text{pure}}$ are normally much larger than 1 (i.e., $\gg 1$ ), $\phi = \frac{N^{\text{free}}_{\text{salt}}}{N^{\text{total}}_{\text{pure}}}$ is the ratio of free water number in salt solution to that of pure water system.

Finally, we have

$$\begin{aligned}
\Delta\Delta S^{\text{water}} &= \Delta S^{\text{water}}_{\text{salt}} - \Delta S^{\text{water}}_{\text{pure}} \\
&= \Delta\Delta S^{\text{translation}} + \Delta\Delta S^{\text{rotation}} + \Delta\Delta S^{\text{vibration}} + \Delta S^{\text{network}} \\
&= \Delta\Delta S^{\text{translation}} + \Delta\Delta S^{\text{rotation}} + \Delta\Delta S^{\text{vibration}} - 4k_{\text{B}} \ln \phi
\end{aligned} \quad \text{(Eq. S13)}$$

where $\Delta\Delta S^{\text{translation}}$ , $\Delta\Delta S^{\text{rotation}}$ , and $\Delta\Delta S^{\text{vibration}}$ can be expressed as follows,

$$\begin{aligned}\Delta\Delta S^{\mathrm{j}} &= \Delta S^{\mathrm{j}}_{\mathrm{salt}} - \Delta S^{\mathrm{j}}_{\mathrm{pure}} \\ &= \left(S^{\mathrm{j\text{-}bound}}_{\mathrm{salt}} - S^{\mathrm{j\text{-}free}}_{\mathrm{salt}}\right) - \left(S^{\mathrm{j\text{-}bound}}_{\mathrm{pure}} - S^{\mathrm{j\text{-}free}}_{\mathrm{pure}}\right) \qquad (\mathrm{Eq.\,S14}) \\ &= S^{\mathrm{j\text{-}free}}_{\mathrm{pure}} - S^{\mathrm{j\text{-}free}}_{\mathrm{salt}}\end{aligned}$$

where j denotes translational or rotational or vibrational components of water entropy, the $S^{\mathrm{j}}$ of bound water molecules by proteins are assumed to be independent of their solution environments and thus cancel out each other. All the parameters in above equation ($\Delta\Delta S^{\mathrm{translation}}$, $\Delta\Delta S^{\mathrm{rotation}}$, $\Delta\Delta S^{\mathrm{vibration}}$, and $\phi$) can be obtained through molecular dynamics simulation and Two-Phase Thermodynamic (2PT) calculation (*33*, *47*).

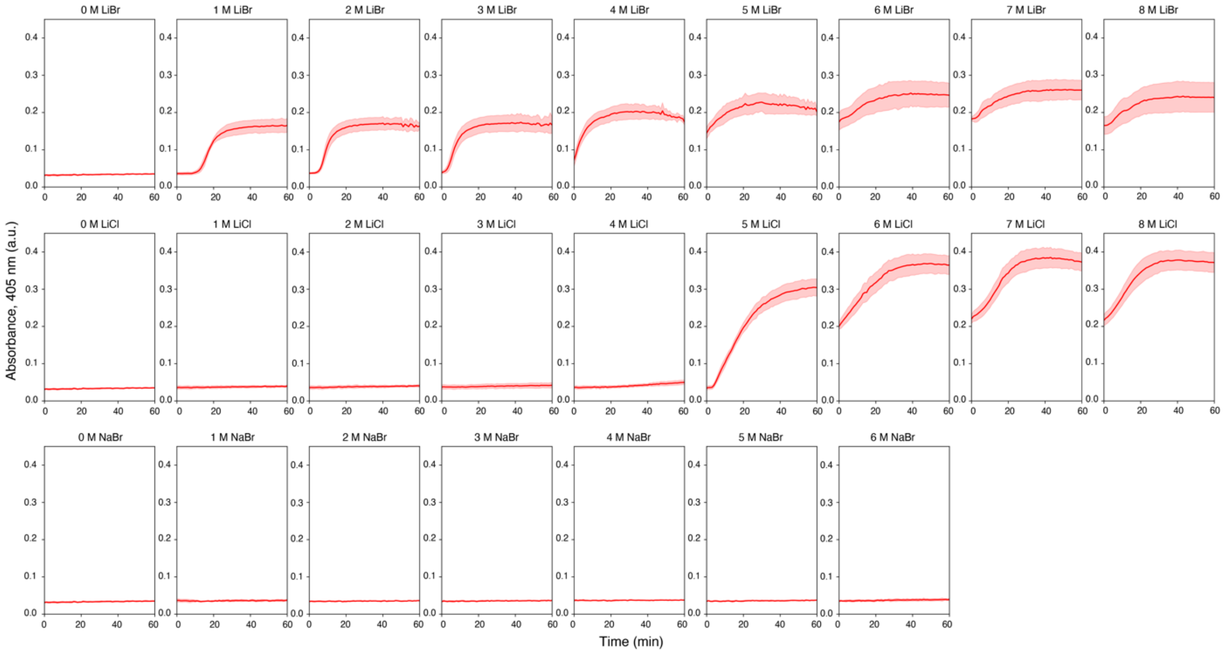


**Fig. S1. Turbidity assay of DHFR in different ions and concentrations.** Absorbance of DHFR in different ion pairs and concentrations at 405 nm wavelength ($OD_{405}$) recorded for 60 minutes. The aggregation propensity consistently follows the trend of LiBr > LiCl > NaBr. Data are presented as mean ± s.d. (n = 4).

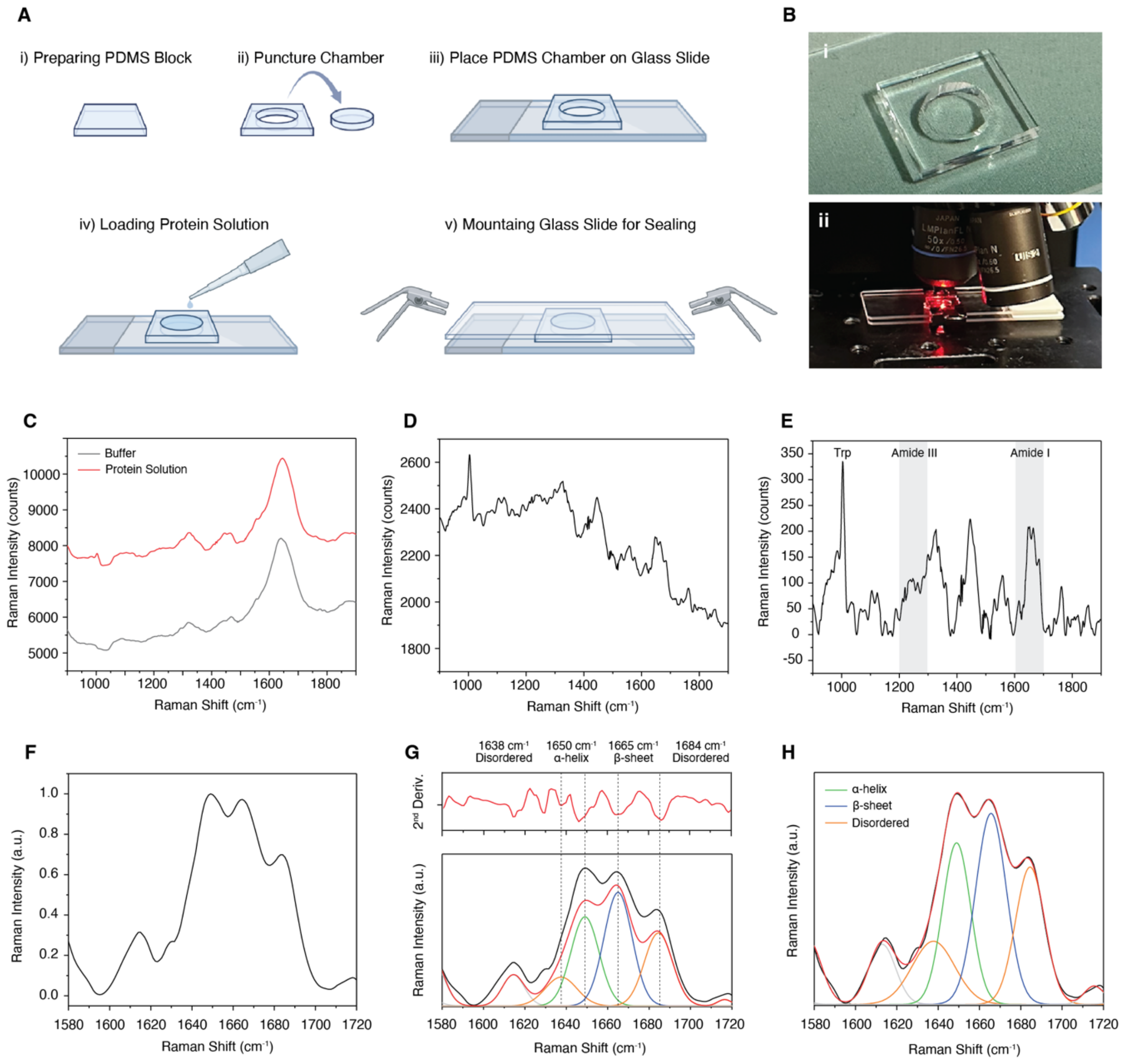


**Fig. S2. Experimental setup and data processing protocol for Raman measurement of DHFR solutions.** (**A**), Schematics of sample preparation for Raman measurement. (**B**), Image of PDMS chamber (i) and the Raman acquisition setup (ii). (**C-H**), Data processing protocol for secondary structure determination of protein sample. (**C**), Collection of buffer background and protein solution under the same condition (e.g., accumulation time). (**D**), Subtracting buffer spectra from protein spectra. (**E**), Subtracting baseline from the previous step, tryptophan peak around 1003 $cm^{-1}$ can be used as reference. (**F**), Smoothing and normalization of amide I band (1600-1700 $cm^{-1}$). (**G**), Using the secondary derivative to locate peaks for deconvolution. (**H**), Peak fitting until convergence, the example is amide I band of DHFR in pure water.

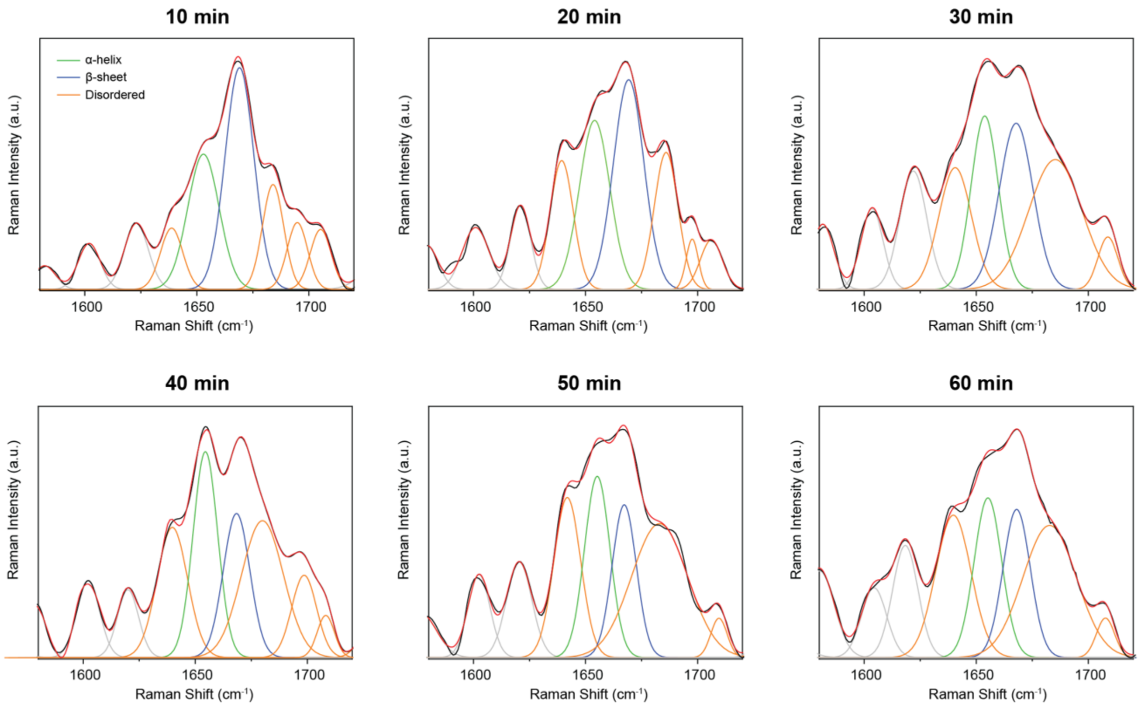


**Fig. S3. Deconvolution of Raman amide I band of DHFR in 2 M LiBr Solution.** DHFR in 2 M LiBr after 10, 20, 30, 40, 50, 60 minutes, respectively. A gradual loss of β-sheet structures around 1665 $cm^{-1}$ and a gradual increase of disordered structures around 1685 $cm^{-1}$ can be observed. Deconvolution of amide I band also suggested no significant change in the percentage of α-helix structure at 1650 $cm^{-1}$.

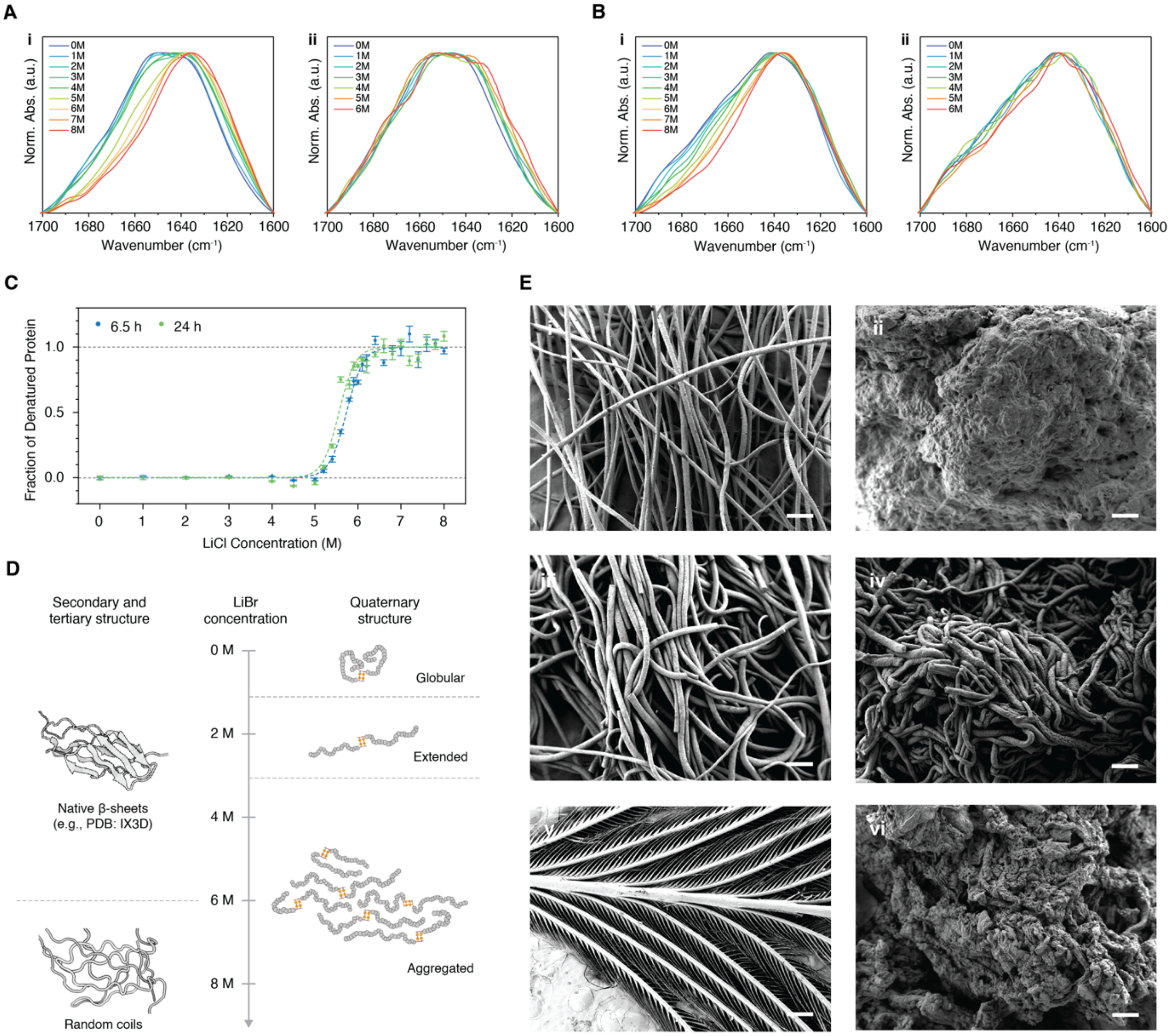


**Fig. S4. LiBr, LiCl, and NaBr exhibit different levels of protein conformational change and denaturation capability.** (**A**), FTIR of the amide I band of DHFR in different concentrations of LiCl (i) and NaBr (ii). (**B**), FTIR of the amide I band of fibronectin in different concentrations of LiCl (i) and NaBr (ii). (**C**), DHFR unfolding curve in LiCl solution measured by tryptophan fluorescence (dots) and fitted by two-state model (dash lines). Data are presented as mean ± s.d. (n = 3). (**D**), Schematics of fibronectin conformational changes under different LiBr concentrations as cross-referenced from FTIR and DLS experiments. Fibronectin conformations start from globular state (0 M), to extended state (~2 M), to aggregation of extended state (4-6 M) until aggregation of denatured state (>6 M). (**E**), SEM image of wool prior (i) and after 48-hour denaturation with 8 M LiBr (ii), 8 M LiCl (iii), and 7 M NaBr (iv). SEM image of feather prior to (v) and after (vi) 48-hour denaturation with 8 M LiBr. Scale bars, 50 µm.

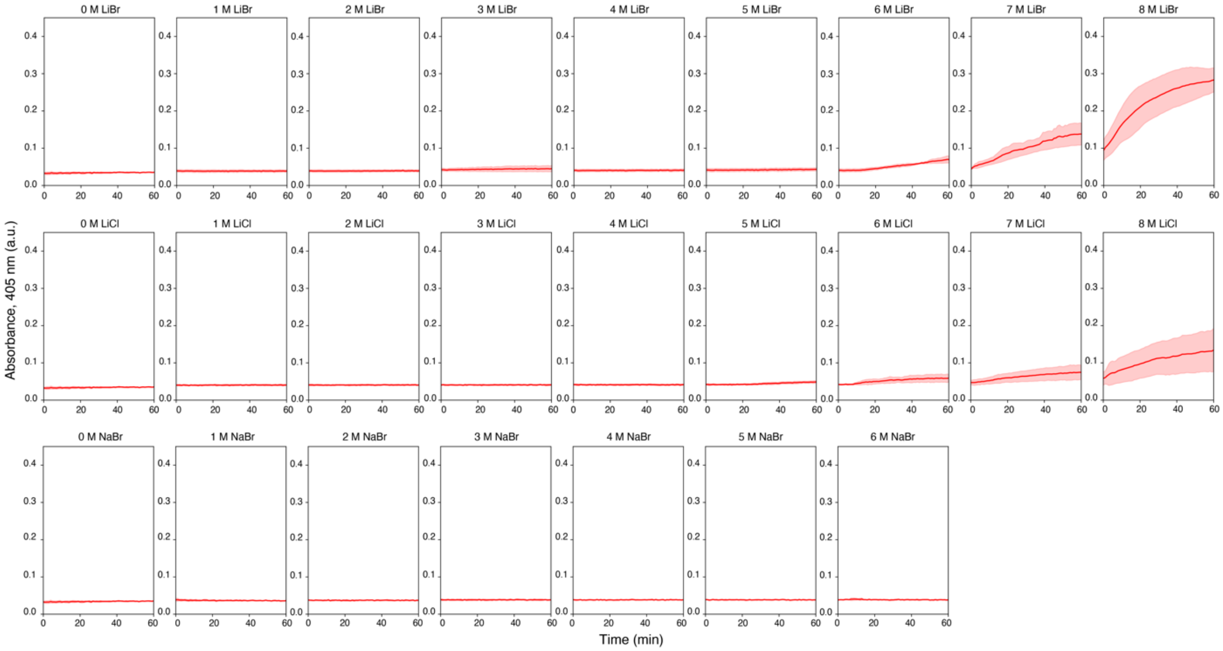


**Fig. S5. Turbidity assay of fibronectin in different ions and concentrations.** Absorbance of fibronectin in different ion pairs and concentrations at 405 nm wavelength ($OD_{405}$) recorded for 60 minutes. Higher concentrations are required to induce significant aggregation compared with DHFR, while the trend of LiBr > LiCl > NaBr remains consistent. Data are presented as mean ± s.d. (n = 4).

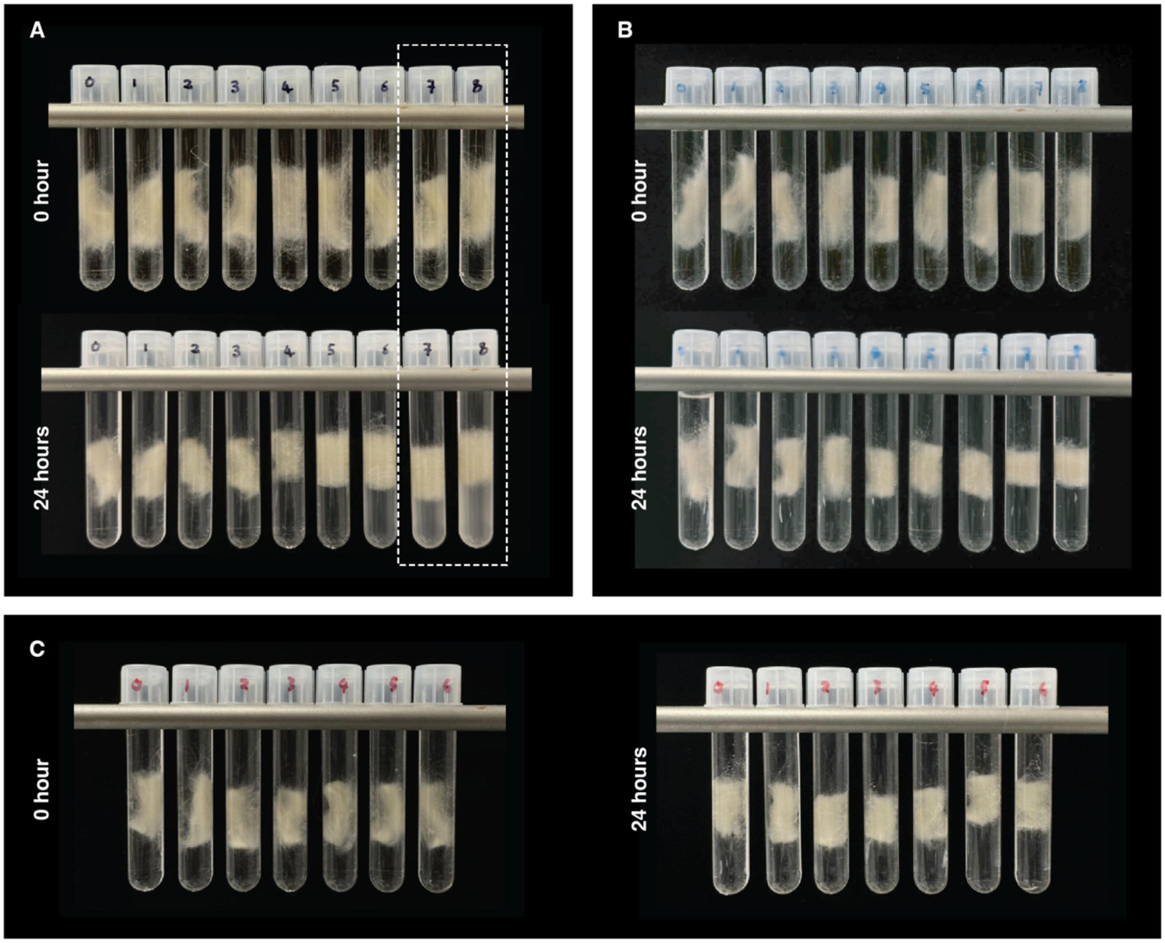


**Fig. S6. Soaking of wool keratin in different ions and concentrations.** Due to the insolubility of keratin in water, turbidity experiments were conducted by soaking keratin in different ion pairs and concentrations for 24 hours at 70 °C to accelerate the kinetics, then measuring the $OD_{405}$ of the solutions. Comparison showed that visible changes only happened at high concentrations of LiBr (**A**), while no difference can be observed in all concentrations of LiCl (**B**) and NaBr (**C**).

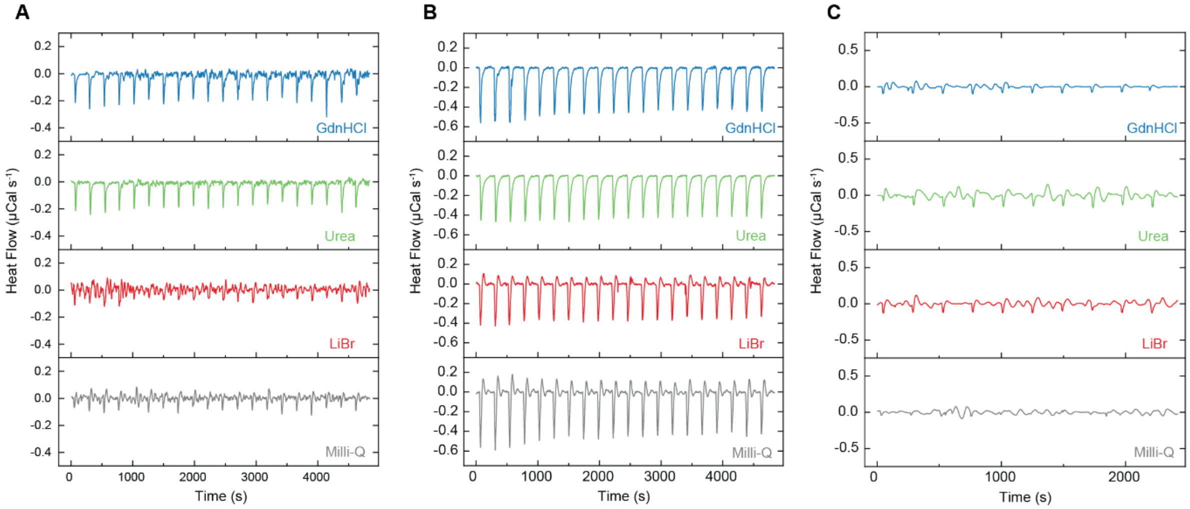


**Fig. S7. Representative curves of ITC experiments.** (**A**), Representative curves of titrating 100 μM of different denaturants into DHFR solution. (**B**), Representative curves of titrating 100 μM of different denaturants into fibronectin solution. Notably, the dilution heat of fibronectin is more significant compared with DHFR potentially due to higher molecular weight. (**C**), Representative curves of titrating 100 μM of different denaturants into Milli-Q water, enthalpy changes were in orders of magnitudes smaller than protein-denaturant interaction. The enthalpy changes of protein-denaturant interaction were calculated by integrating each peak followed by subtraction of dilution heat of protein.

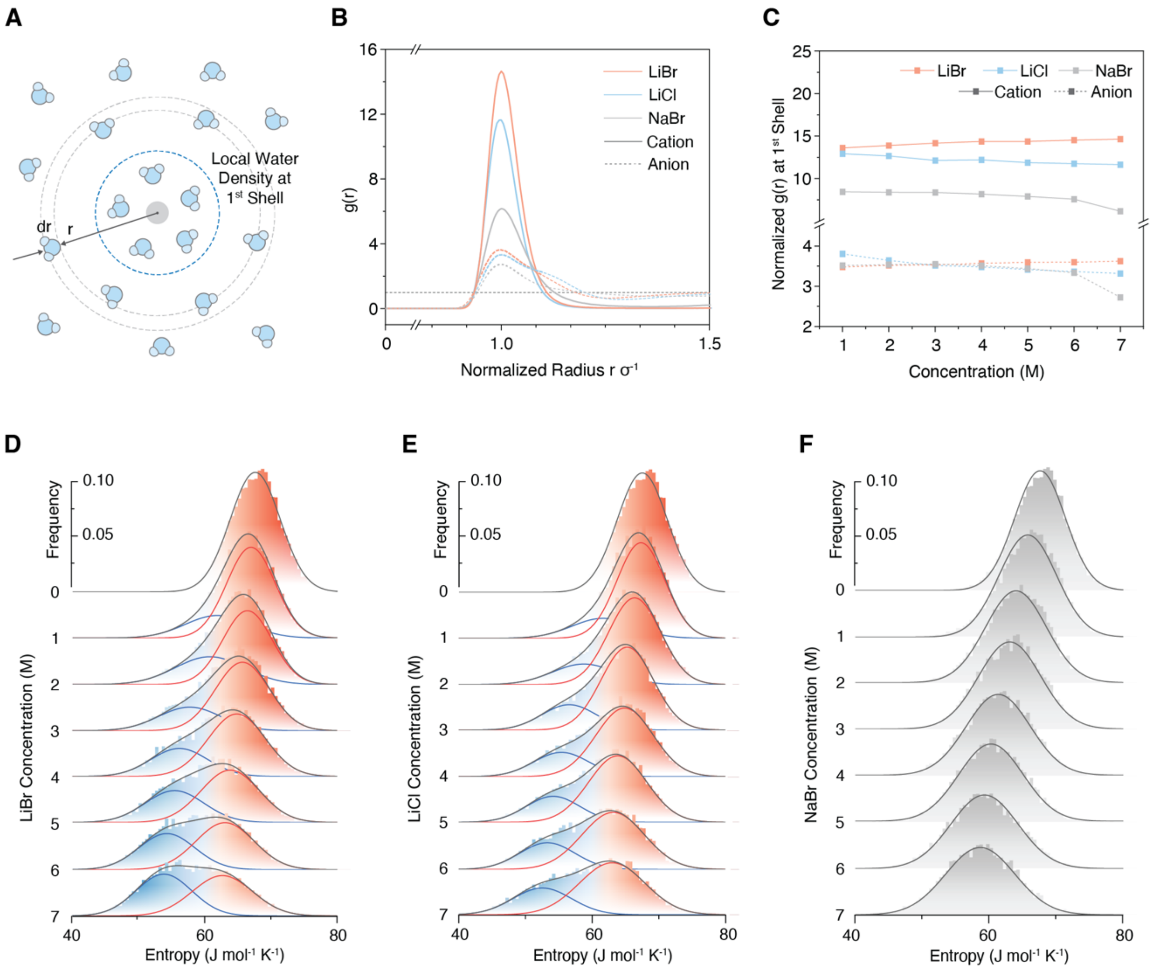


**Fig. S8. Local and global effects of ions on water dynamics.** (**A**), Schematics of the radial distribution function *g(r)* and the local water density within the first hydration shell. (**B**), *g(r)* of oxygen atoms of $H_2O$ molecules around respective cations (solid) and anions (dashed) in 7 M LiBr, LiCl, and NaBr solution. (**C**), Normalized *g(r)* of oxygen atoms of $H_2O$ molecules at first hydration shell around respective cations (solid) and anions (dashed) in different concentrations of LiBr, LiCl, and NaBr. (**D-F**), Water molecular entropy distribution in LiBr, LiCl, and NaBr, respectively, at different concentrations.

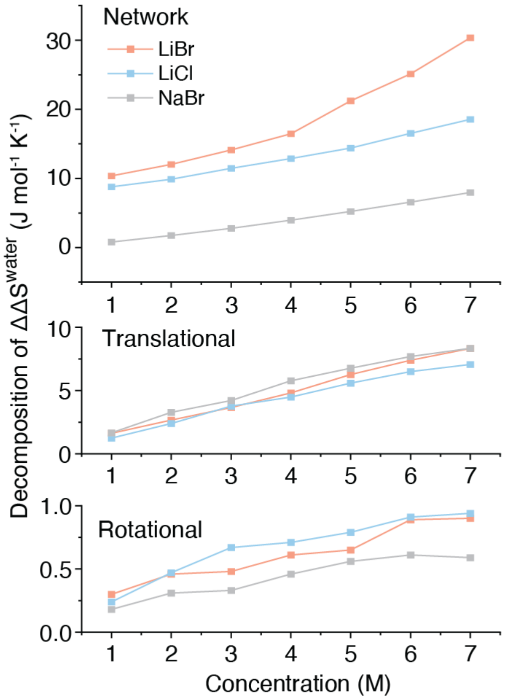


**Fig. S9. Decomposition of the total water entropy penalty decrease.** Decomposition of the total water entropy penalty decrease presented in Fig. 2D. The contribution of network entropy penalty appears to be the predominant factor over rotational and translational entropy of individual water molecules.

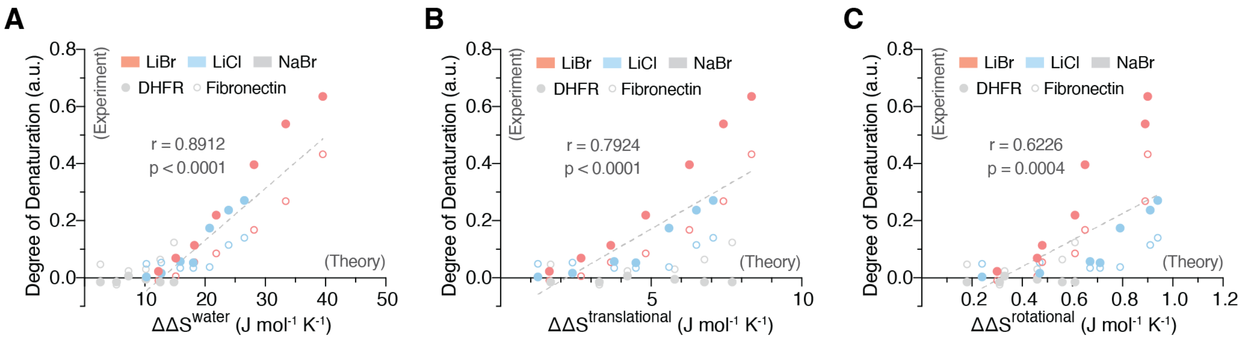


**Fig. S10. Correlation between the theoretical value of entropy penalty decrease and the protein denaturation behavior observed from FTIR experiments. (A)**, Correlation between the total water entropy penalty decrease and denaturation behavior. The high Pearson's r value (0.8912) indicates a strong correlation. (**B**), Correlation between the translational entropy penalty decrease and denaturation behavior. (**C**), Correlation between the rotational entropy penalty decrease and denaturation behavior. Compared to the weaker correlations with rotational and translational entropy penalties, the decrease in network entropy penalty emerges as the predominant factor.

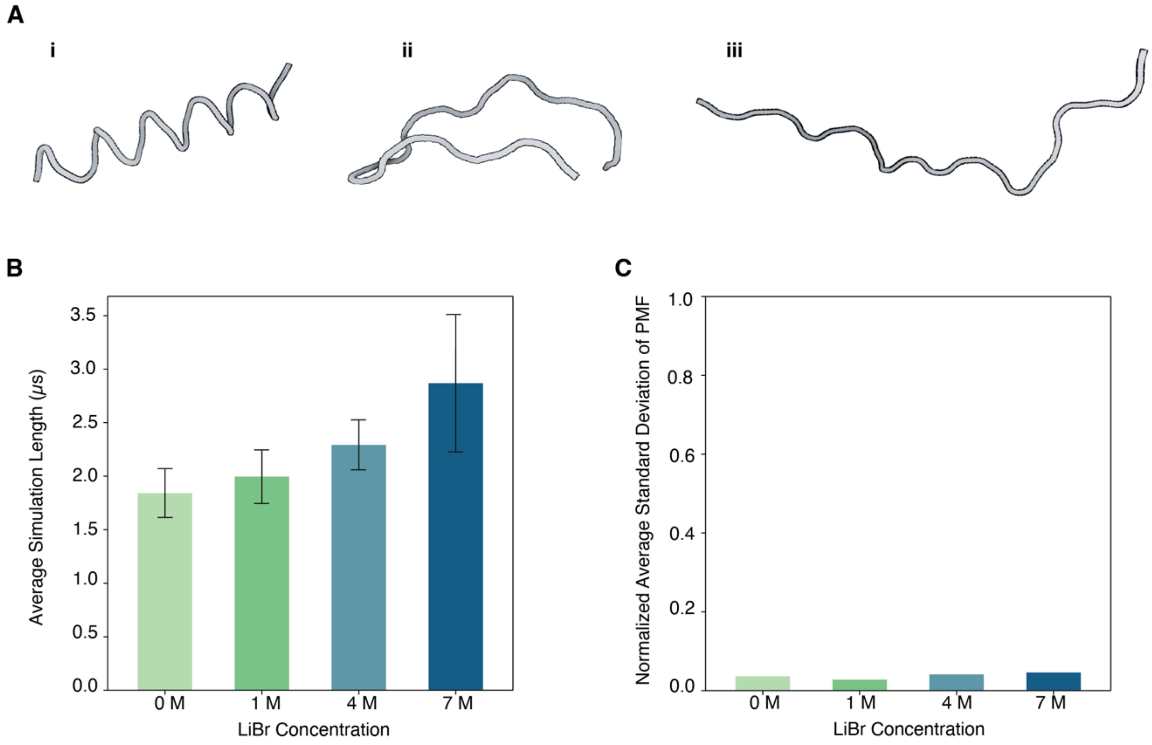


**Fig. S11. Metadynamics simulations initial configurations and results summary.** (**A**), Three different initial configurations for metadynamics simulations (i-iii are structure 1-3, respectively, as denoted in Fig. S12-15). (**B**), Simulation trajectory length summary of metadynamics simulations with different LiBr concentrations. Data are presented as mean ± s.d. (n = 6). (**C**), Normalized average standard deviation of Potential of Mean Force (PMF) from metadynamics simulations for different LiBr concentrations.

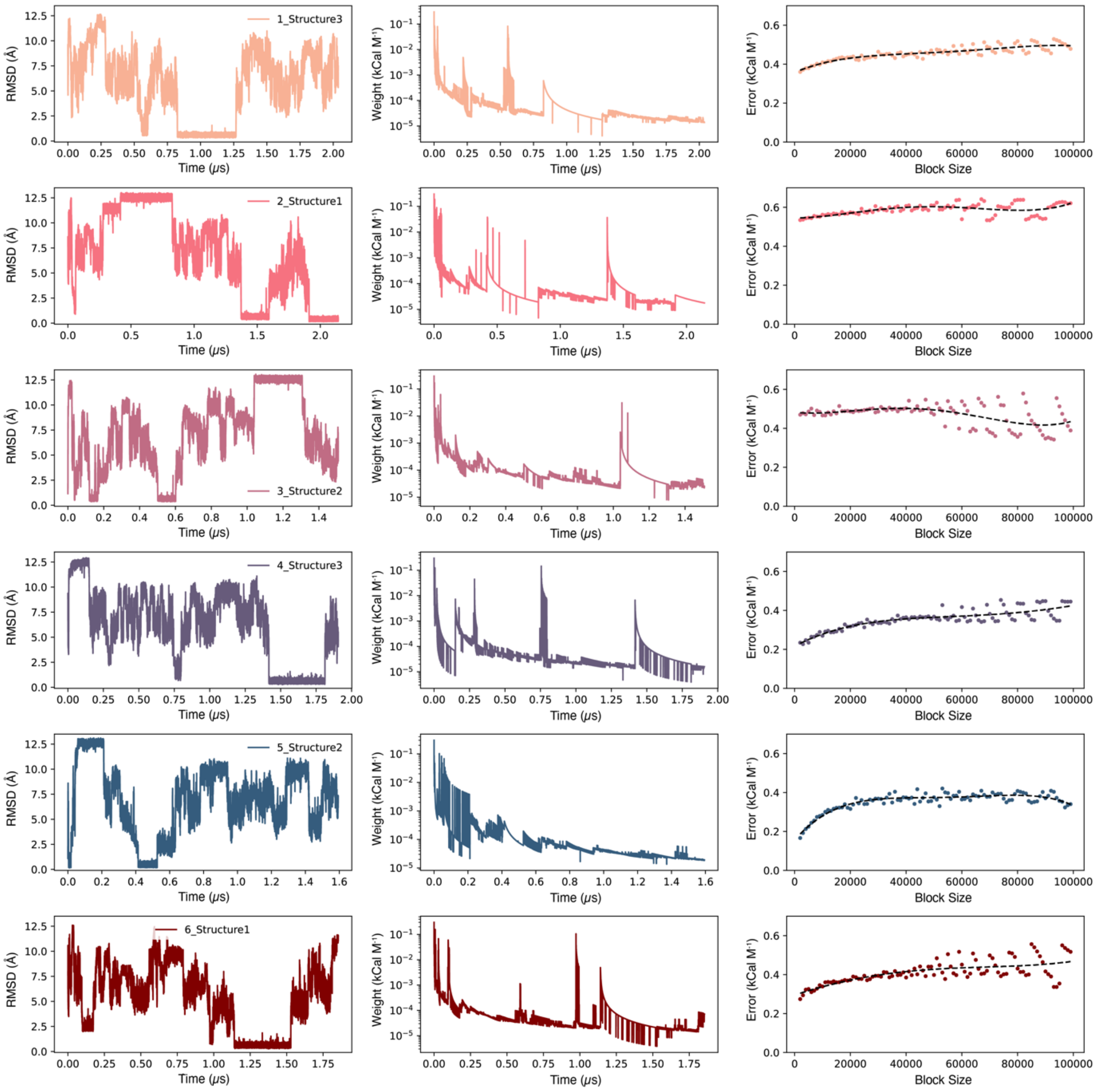


**Fig. S12. Convergence confirmation of metadynamics simulations for 0 M LiBr.** Rigorous criteria have been implemented to ensure the accuracy of free energy landscapes derived from metadynamics simulations. Initially, simulations are conducted until the collective variable (CV) iterates over its entire possible range, in this case, 0 to 12.5 Å (first column). Secondly, the final height of the Gaussian bias potential added at each CV position must be less than $5*10^{-3}$ kCal $M^{-1}$ (second column). Lastly, the convergence of the simulations is further confirmed by observing a plateau in the average standard deviation of the potential of mean force (PMF) as calculated from block analysis, with increasing block sizes (third column). Six converged simulations out of nine for each LiBr concentration were used for the final protein free energy landscape construction as shown here.

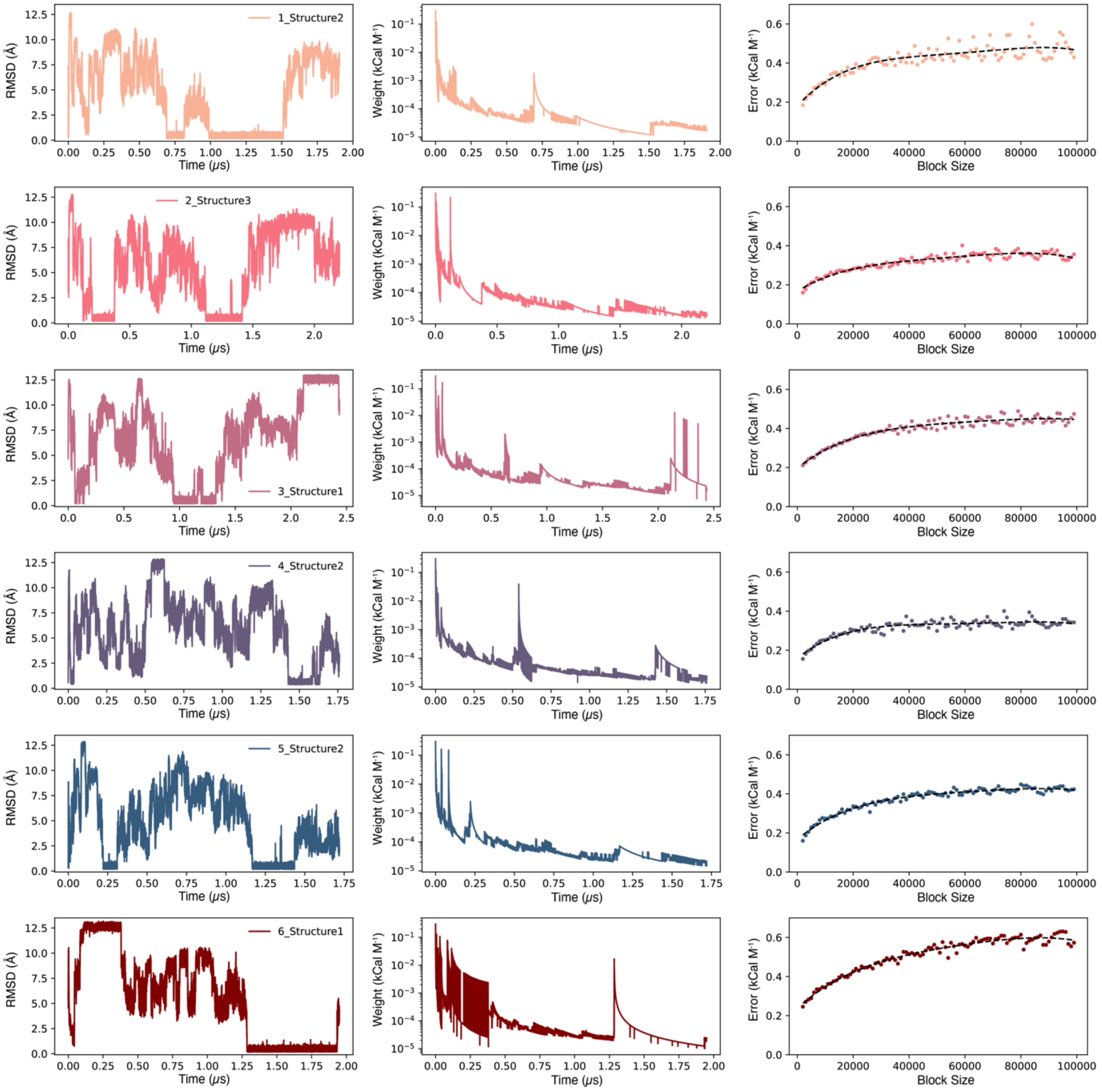


**Fig. S13. Convergence confirmation of metadynamics simulations for 1 M LiBr.** Rigorous criteria have been implemented to ensure the accuracy of free energy landscapes derived from metadynamics simulations. Initially, simulations are conducted until the collective variable (CV) iterates over its entire possible range, in this case, 0 to 12.5 Å (first column). Secondly, the final height of the Gaussian bias potential added at each CV position must be less than $5*10^{-3}$ kCal $M^{-1}$ (second column). Lastly, the convergence of the simulations is further confirmed by observing a plateau in the average standard deviation of the potential of mean force (PMF) as calculated from block analysis, with increasing block sizes (third column). Six converged simulations out of nine for each LiBr concentration were used for the final protein free energy landscape construction as shown here.

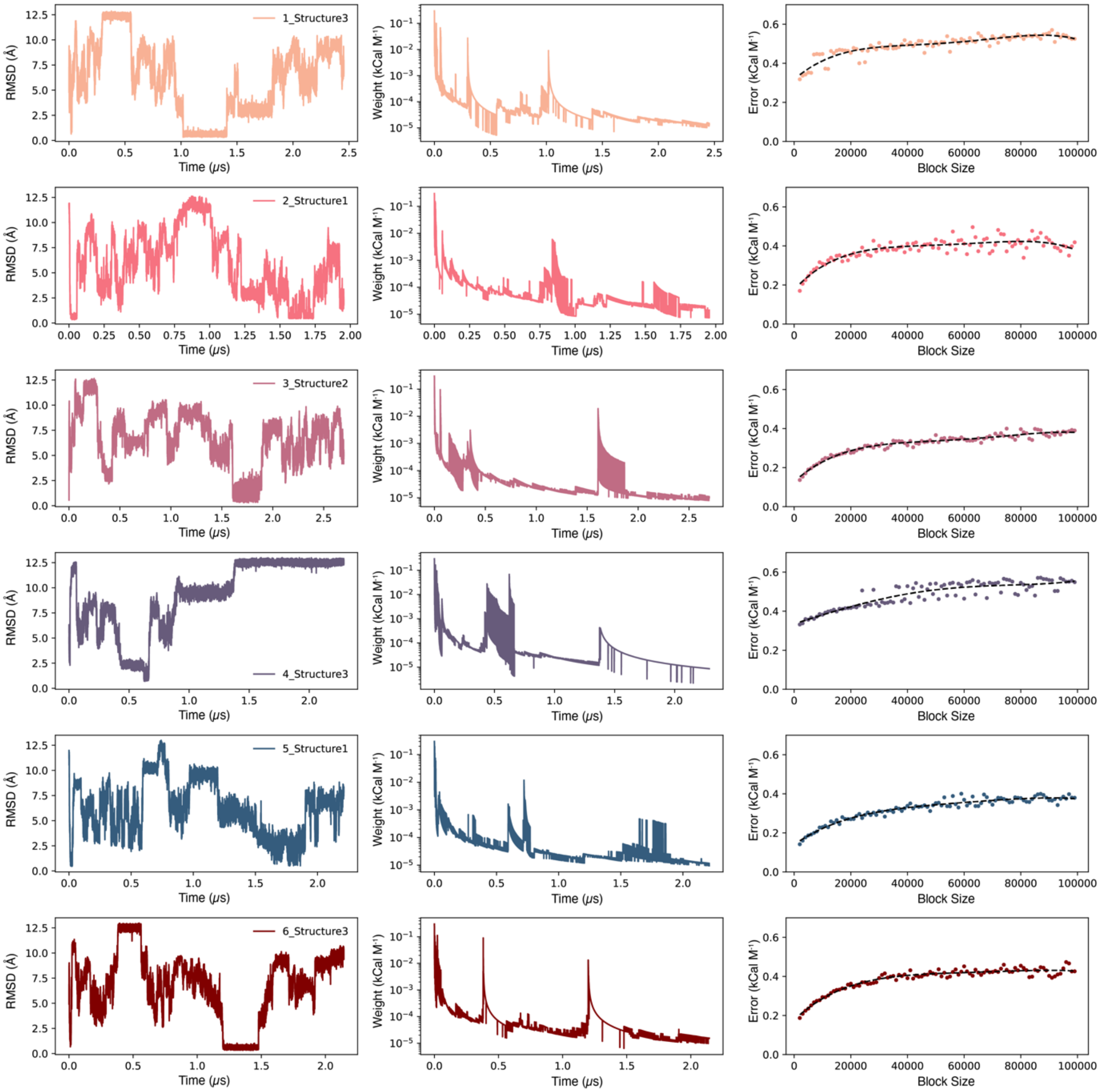

**Fig. S14. Convergence confirmation of metadynamics simulations for 4 M LiBr.** Rigorous criteria have been implemented to ensure the accuracy of free energy landscapes derived from metadynamics simulations. Initially, simulations are conducted until the collective variable (CV) iterates over its entire possible range, in this case, 0 to 12.5 Å (first column). Secondly, the final height of the Gaussian bias potential added at each CV position must be less than $5*10^{-3}$ kCal $M^{-1}$ (second column). Lastly, the convergence of the simulations is further confirmed by observing a plateau in the average standard deviation of the potential of mean force (PMF) as calculated from block analysis, with increasing block sizes (third column). Six converged simulations out of nine for each LiBr concentration were used for the final protein free energy landscape construction as shown here.

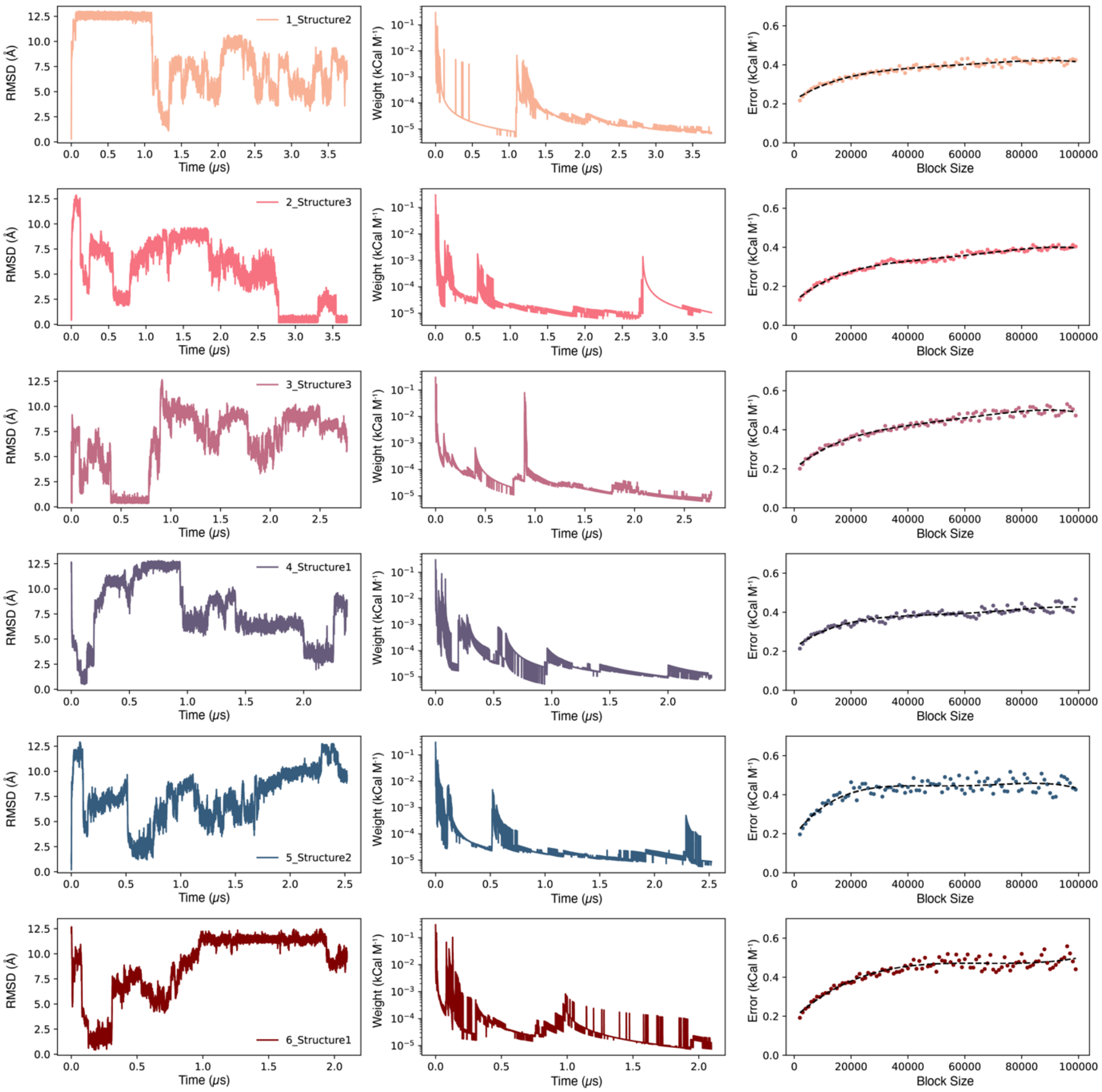


**Fig. S15. Convergence confirmation of metadynamics simulations for 7 M LiBr.** Rigorous criteria have been implemented to ensure the accuracy of free energy landscapes derived from metadynamics simulations. Initially, simulations are conducted until the collective variable (CV) iterates over its entire possible range, in this case, 0 to 12.5 Å (first column). Secondly, the final height of the Gaussian bias potential added at each CV position must be less than $5*10^{-3}$ kCal $M^{-1}$ (second column). Lastly, the convergence of the simulations is further confirmed by observing a plateau in the average standard deviation of the potential of mean force (PMF) as calculated from block analysis, with increasing block sizes (third column). Six converged simulations out of nine for each LiBr concentration were used for the final protein free energy landscape construction as shown here.

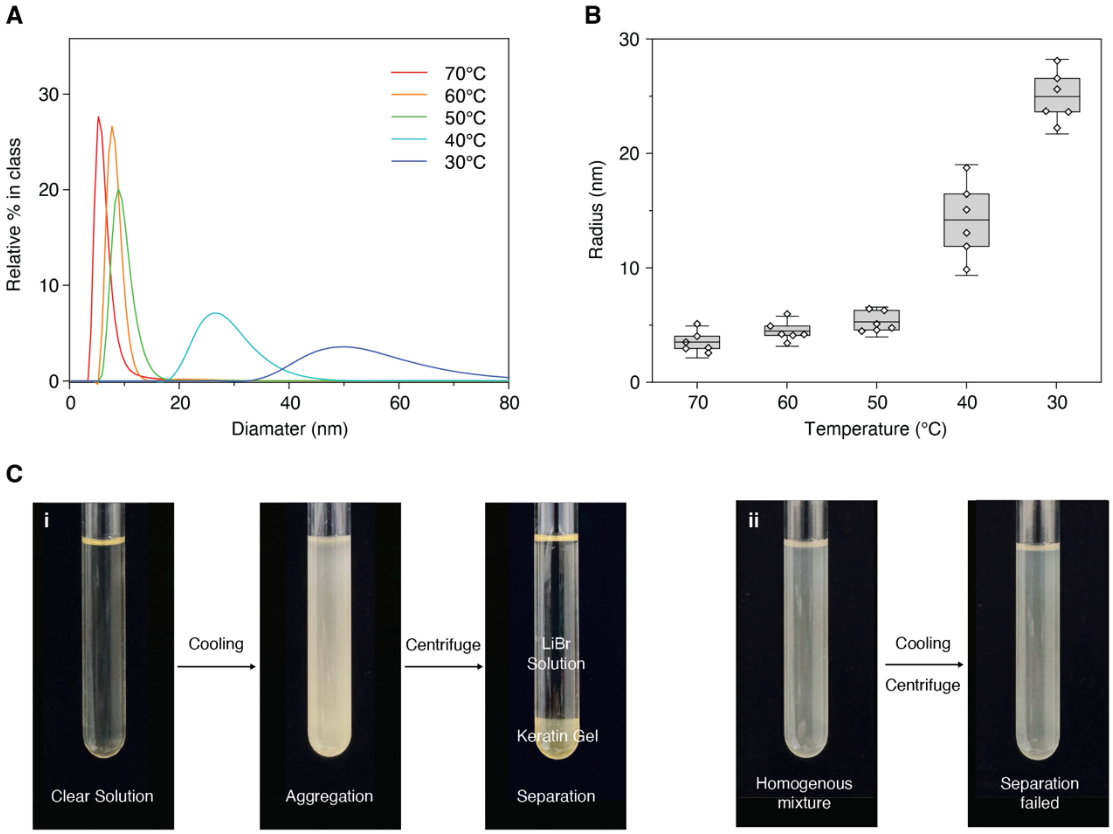


**Fig. S16. Spontaneous aggregation and separation of keratin extracted with LiBr solution.** (**A** and **B**), Size distribution of denatured keratin in 8 M LiBr solution suggests a spontaneous aggregation as temperature decrease from DLS. (**C**), Keratin solution extracted with LiBr undergoes spontaneous aggregation upon cooling and can be easily separated into condensed gel after centrifuging (i). Keratin solution extracted with urea remains homogenous after the same process, requiring dialysis for separation (ii).

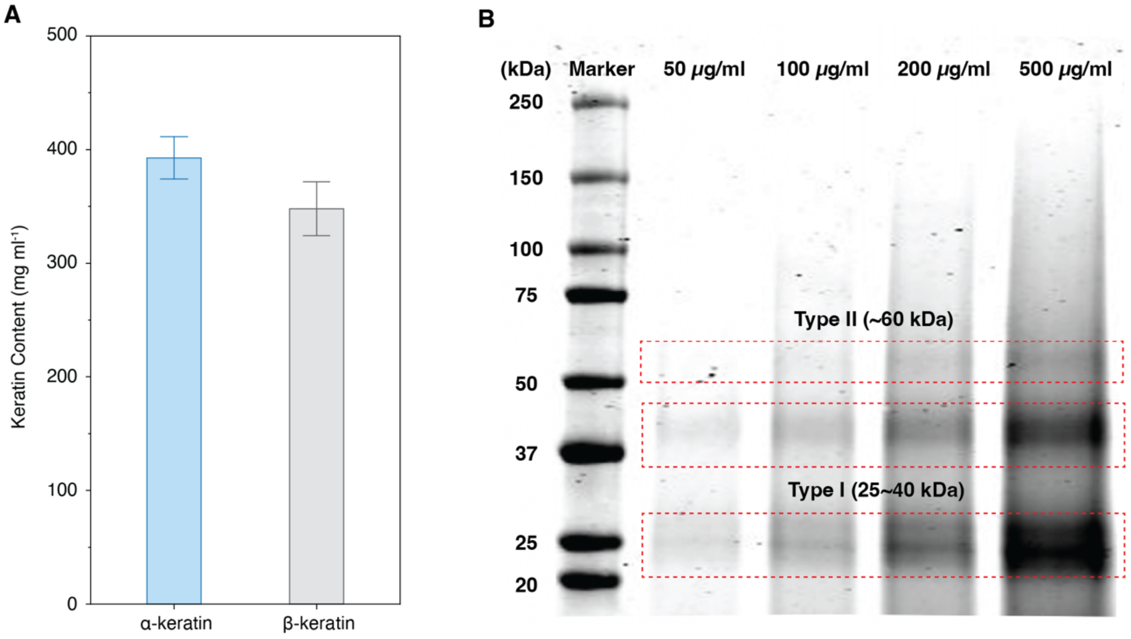


**Fig. S17. Keratin content of extracted gel and confirmation of keratin composition.** (**A**), Keratin content of aggregated gel after separation. Data are presented as mean ± s.d. (n = 6). (**B**), SDS-Page of extracted keratin contains type I keratin with a molecular weight around 60 kDa and type II keratin with a molecular weight around 25~40 kDa. The presence of additional smeared bands at higher Mw may be attributed to connected dimers or tetramers, while the smear below 20 kDa may be attributed to other matrix proteins and helical fragments within the wool.

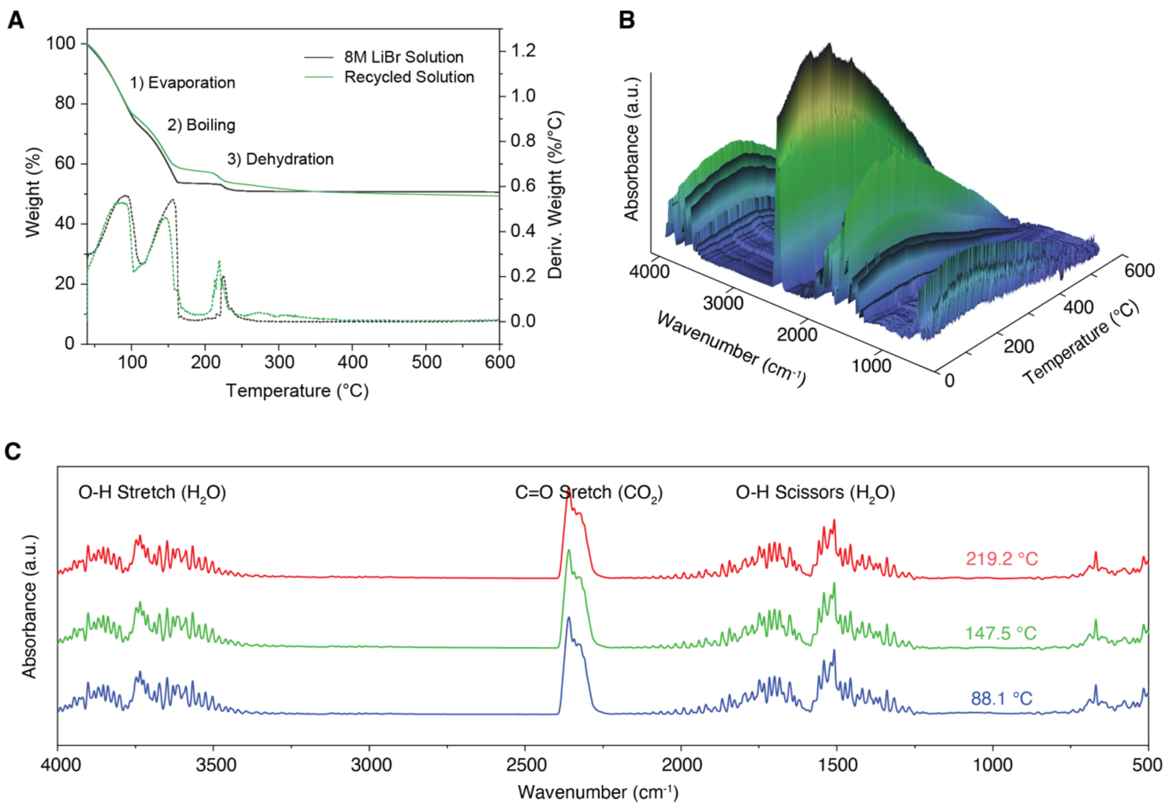


**Fig. S18. Composition analysis of the LiBr solution after close-loop recycling (cycle 5).** (**A**), Comparison of the TGA profile shows no major change in the LiBr solution after 5 cycles of extraction. (**B** and **C**), FTIR of the evolving gas during TGA measurement of recycled LiBr solution. Despite the intensity change due to different evaporation rate at different stages, the chemical composition showed little variation throughout the heating process, indicating a stable feed of water vapor.

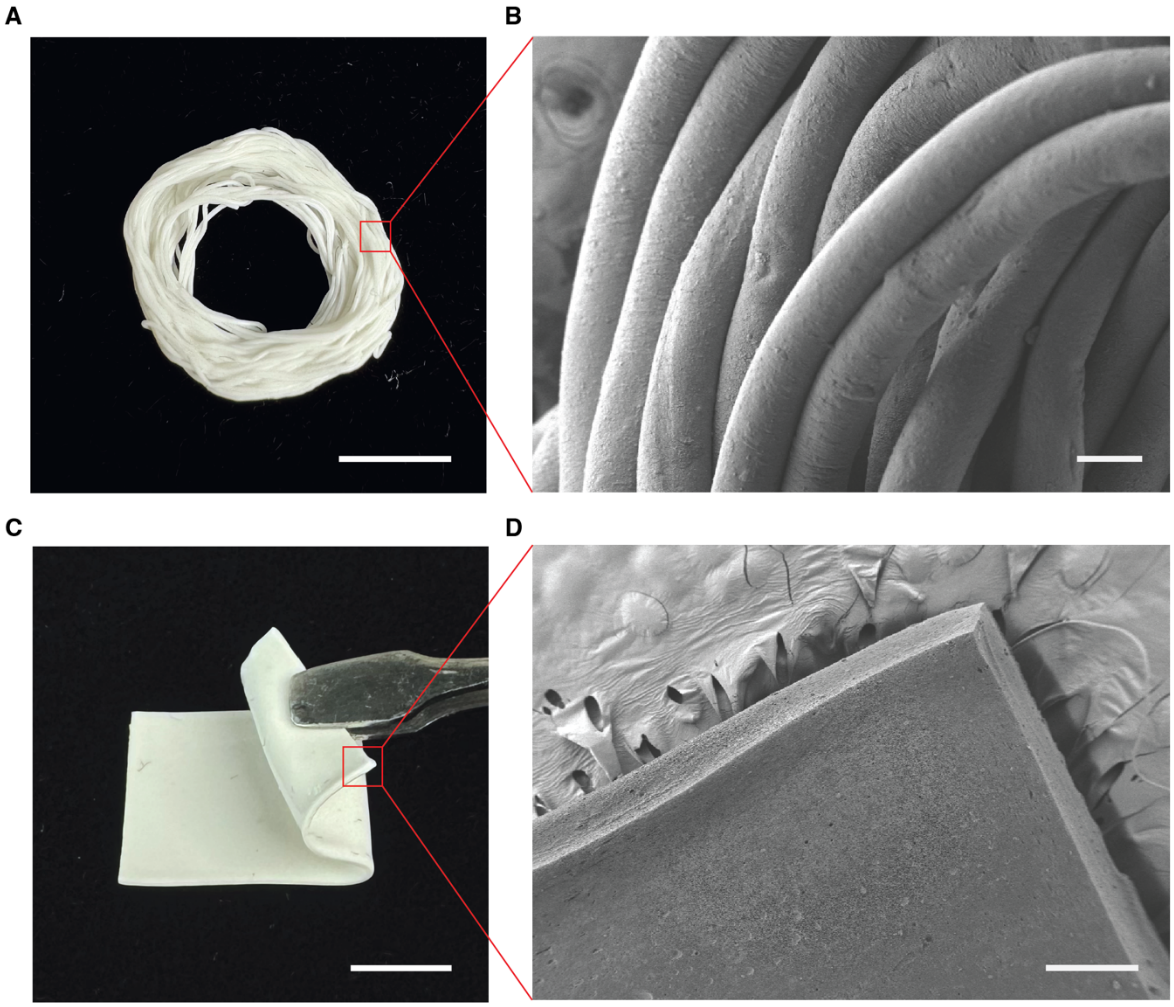


**Fig. S19. SEM images of regenerated keratin fiber and membrane.** (**A** and **B**), Optical and SEM images of keratin fibers fabricated by injecting keratin gel into $NaH_2PO_4$ solution. Scale bars, 1 cm for optical image, 100 μm for SEM image. (**C** and **D**), Optical and SEM images of keratin membrane casted with keratin gel. Cross-section and edge were created with razor blades. Scale bars, 1 cm for optical image, 300 μm for SEM image.

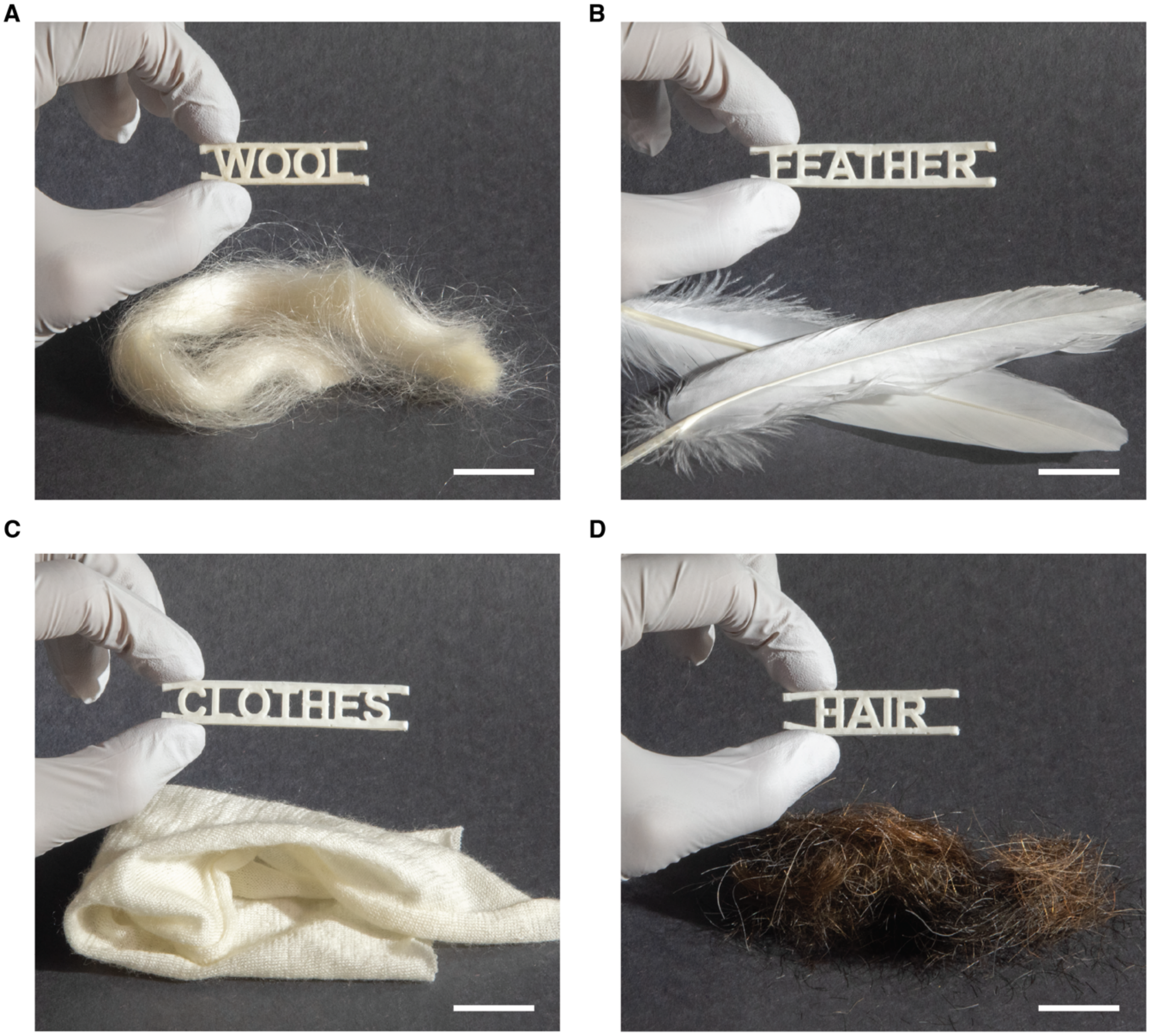


**Fig. S20. Regenerated keratin materials posted next to their original sources.** Different keratin samples were regenerated from (**A**) wool, (**B**) goose feathers, (**C**) wool clothes, and (**D**) human hair. Keratin samples were shaped into the word describing their sources using the injection molding method. Scale bars, 1 cm.

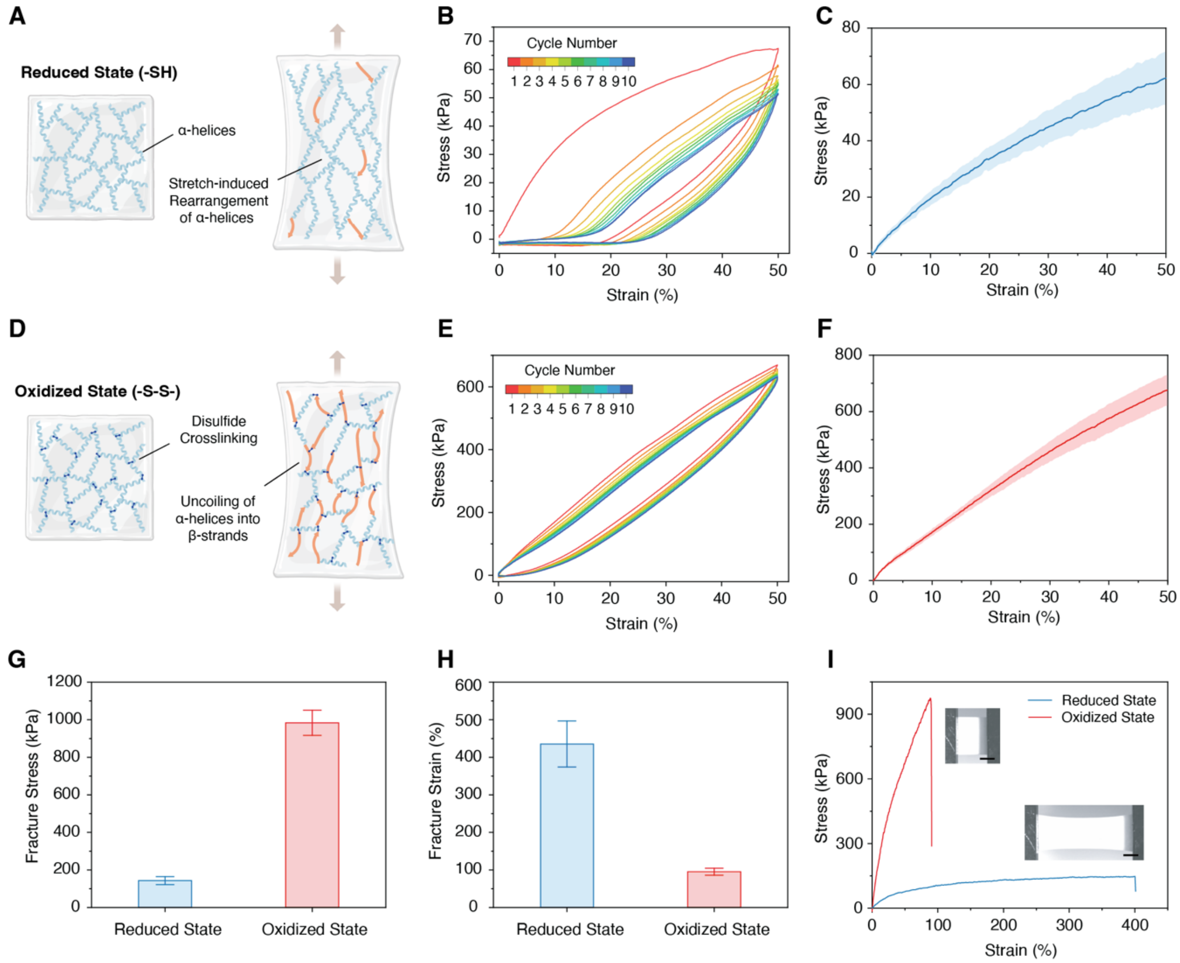


**Fig. S21. Tunable mechanical properties of regenerated keratin material.** (**A**), Schematic of the α-keratin in reduced state and directional rearrangement of α-helices induced by stretching. (**B**), Cyclic loading of reduced state keratin to 50% strain. (**C**), Stress-strain curve of reduced state keratin. Data are presented as mean ± s.d. (n = 4). (**D**), Schematic of the α-keratin in oxidized state and uncoiling of helices upon stretch. (**E**), Cyclic loading of oxidized state keratin to 50% strain. (**F**), Stress-strain curve of oxidized state keratin. Data are presented as mean ± s.d. (n = 4). (**G** and **H**), Fracture stress and fracture strain of regenerated keratin in reduced state and oxidized state. Data are presented as mean ± s.d. (n = 6). (**I**), Comparison of stress-strain curves between reduced and oxidized keratin. Scale bars, 2 mm.

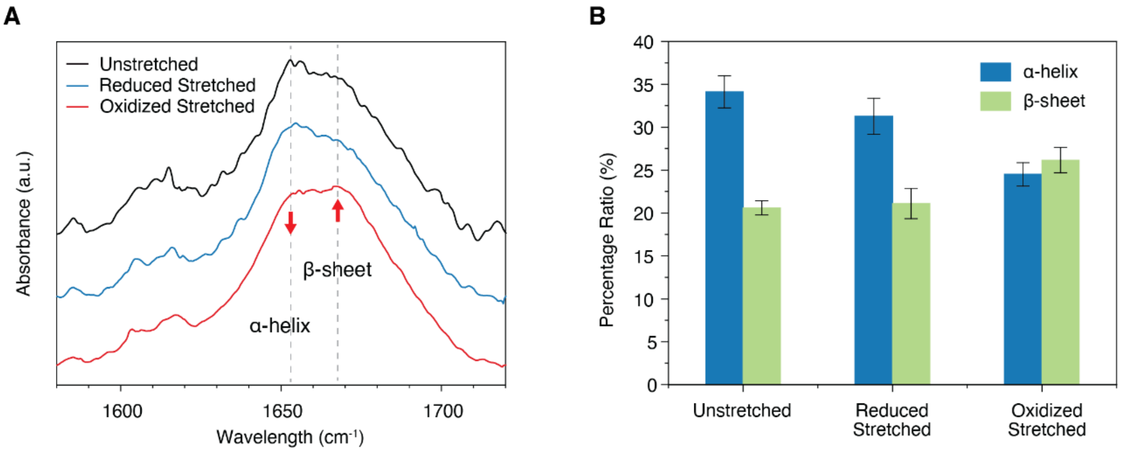


**Fig. S22. Structural behavior of regenerated keratin under stress.** (**A**), Comparison of the amide I band of unstretched keratin, reduced keratin under 50% strain and oxidized keratin under 50% strain. (**B**), Change of secondary structure percentage under 50% strain quantified from Raman deconvolution of amide I band presented in (**A**). A more significant decrease of α-helix can be observed within the oxidized sample, indicating the uncoiling of helices into metastable β-sheets (n = 3).

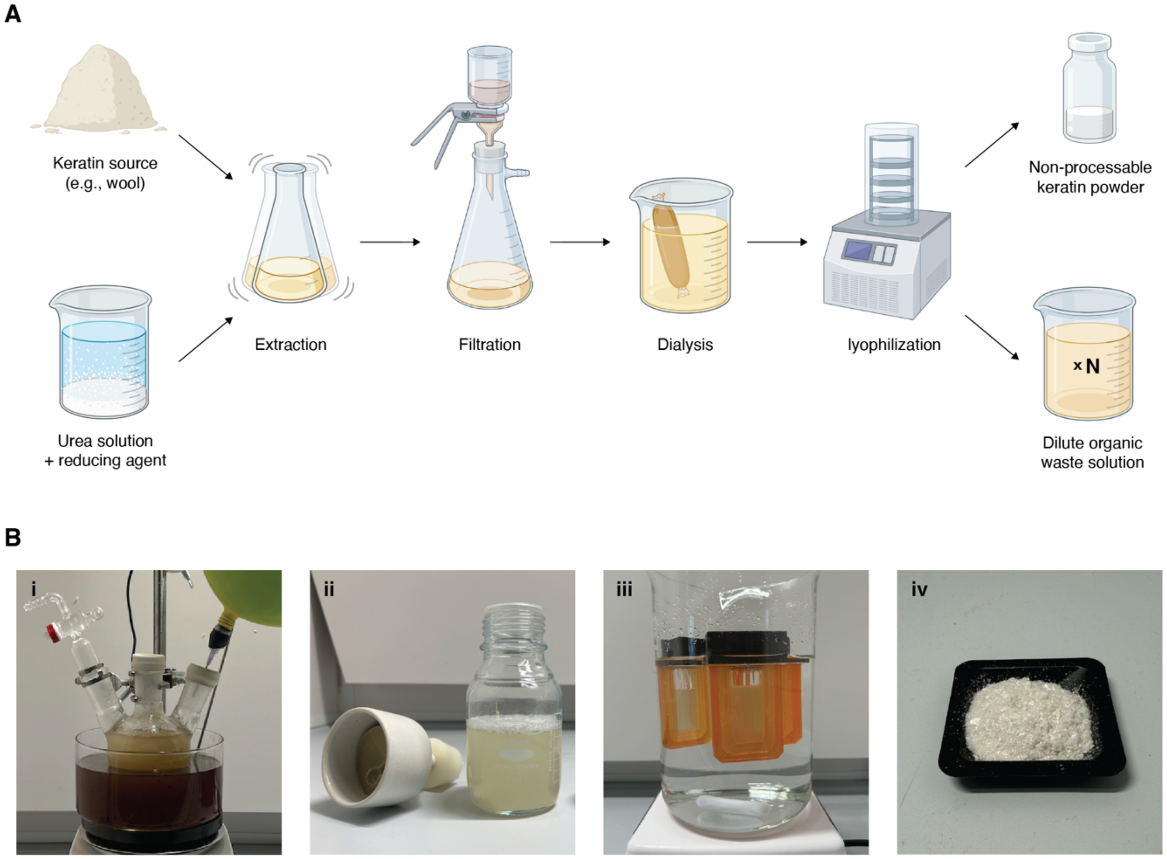


**Fig. S23. Traditional extraction process of keratin using organic denaturing agents.** (**A**), Flowchart of keratin extraction with organic denaturants (e.g., urea). A final product of non-processable keratin powder can be obtained with a large volume of dilute organic waste solution. (**B**), Images of the extraction process steps including extraction (i), filtration (ii), dialysis (iii), and post-lyophilization keratin powder (iv). The lyophilized keratin powder is insoluble and remains dispersed, requiring blending with another material for further processing. Due to its inability to form a continuous structure, the shape-memory effect of keratin could not be observed.

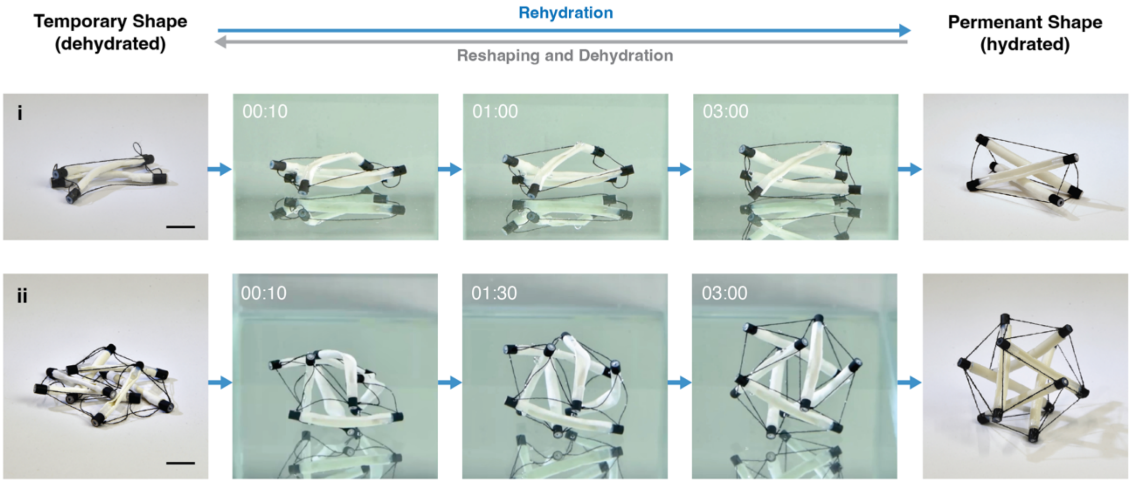


**Fig. S24. Shape-memory effect of regenerated keratin as tensegrity structures.** Dehydrated tensegrity structures spontaneously recover to their original conformations upon rehydration within a timescale of minutes, demonstrated with a three-rod tensegrity structure (i) and a six-rod tensegrity structure (ii). Full recovery observed in both samples regardless of complexity difference. Scale bars: 1.5 cm.

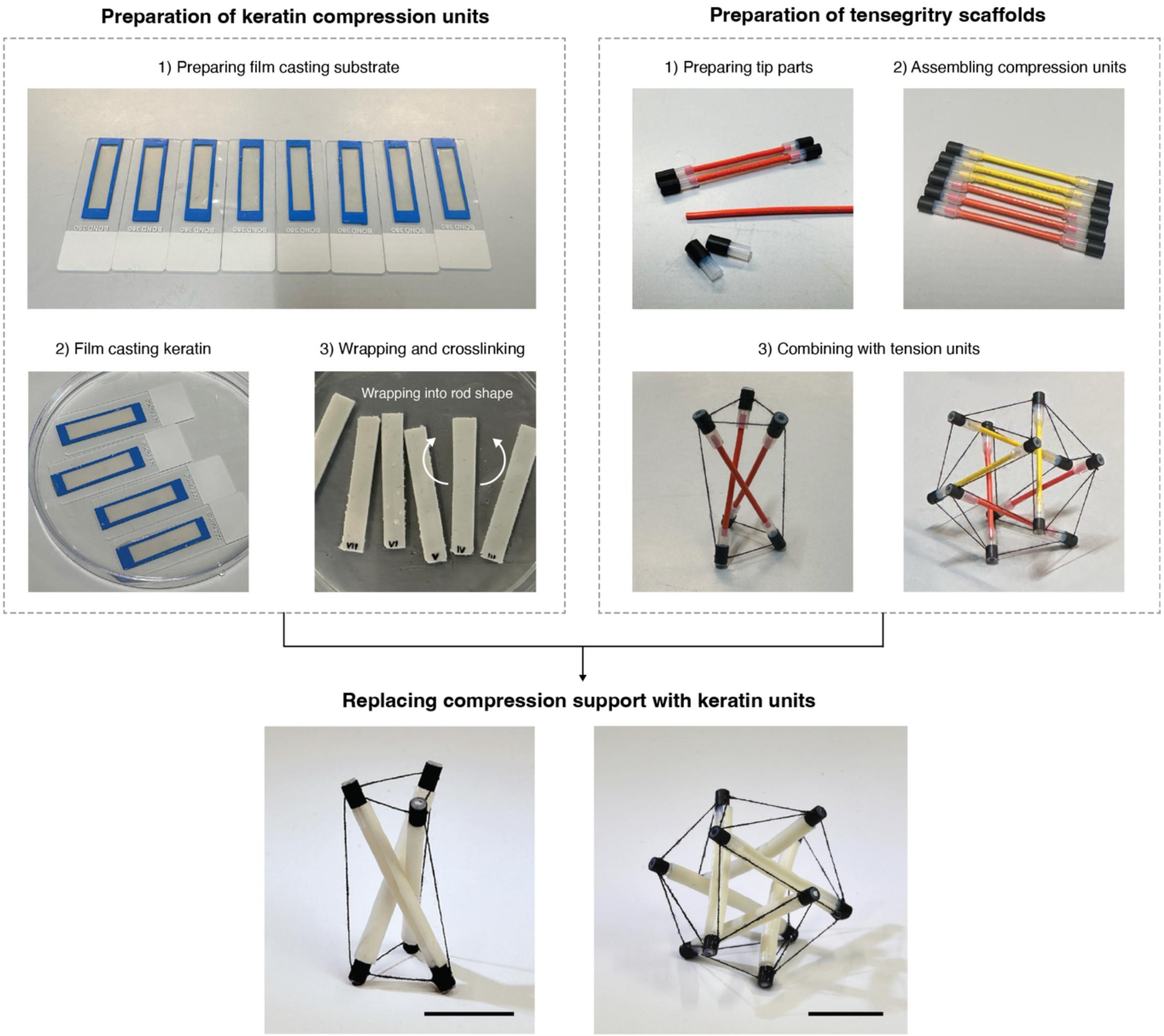


**Fig. S25. Fabrication process of tensegrity structures with keratin.** Three-rod and six-rod tensegrity structures were fabricated with compression units made of crosslinked keratin. Compression units were prepared by film casting keratin into rectangle sheets, then wrapped into rod shapes to provide stronger structural support. Tensegrity scaffolds were first built with temporary compression units assembled with rubber tips. Nylon strings were threaded through the tips and tightened to form tension units. After the structures were adjusted to stabilization, keratin rods were glued onto the tips with cyanoacrylate glue, followed by the removal of temporary compression support. Both tensegrity structures remain stable in a hydrated environment. Scale bars: 2 cm.

**Table S1. Summary of current keratin extraction strategies**

| Method | Example Procedure and Result | Keratin Source | Reference |
|---|---|---|---|
| Organic denaturant | 8 M guanidine hydrochloride (GdnHCl) and 1.66 M 2-mercaptoethanol at 60-70 °C for 18 h, then dialyzed. 41.3% yield. Type I and type II keratin identified | Wool | Ozaki et al. (*51*) |
| | 7 M urea, 2 M thiourea, 50 mM dithiothreitol (DTT), and 50 mM Tris (pH 9.3) for 18 h, then dialyzed and lyophilized. 37 types of peptides identified | Wool | Choudhury et al. (*52*) |
| | 8 M urea, 0.5 M sodium metabisulfite ($Na_2S_2O_5$), 0.05 M SDS, and 5 N NaOH (pH 6.5) at 65 °C for 3 h, filtered and dialyzed against distilled water for 3 days. α-helix structures recovered | Wool | Kadirvelu et al. (*53*) |
| | 1) 8 M urea; 0.26 M SDS, 1.66 M 2-mercaptoethanol at 50 °C for 12 h, 17% yield. 2) 8 M urea, 0.165 M L-cysteine, and 3M NaOH (pH 10.5) at 75 °C for 5 h, 26% yield. Both require dialysis for 3 days | Wool | Ramya et al. (*54*) |
| Acidic/basic/ oxidative hydrolysis | 2% peracetic acid for 30 h, followed by 0.2 N ammonia and precipitation with 2 N HCl | Wool | Earland et al. (*55*) |
| | 0.5 N NaOH (pH 13.9) at 62-65°C for 3h, dialyzed and lyophilized, Mw around 6.5 kDa | Wool | Cardamone (*56*) |
| | 2% NaOH at 80 °C for 3 h, 25% yield, Mw of proteins less than 20 kDa | Human hair | Zhang et al. (*57*) |
| | 0.125 M sodium sulfide ($Na_2S$) at 40 °C for 4 h, filtration and dialysis for 3 days, 10% yield of polypeptides with no secondary structures | Wool | Ramya et al. (*54*) |
| | 10 g/L $Na_2S$ solution in $N_2$ atmosphere at 30 °C for 1 h, Mw around 10 kDa (70% product) | Feather | Poole et al. (*58*) |
| | KOH: NaOH (14:1) at 0.5 to 3% (w/v) for 24 h, facilitated by ultrasonic irradiation for 30 min at 20 kHz (40% amplitude of 750 W) | Human hair | Bhat et al. (*59*) |

| | | | |
|---|---|---|---|
| Ionic liquid (IL) | 1-butyl-3-methylimidazolium [Bmim]Cl, [Bmim]Br, [Bmim]$BF_4$, and [Bmim]$PF_6$ at 100-130°C for 10-24 h. Disappearance of α-helix structures reported | Wool | Xie et al. (*60*) |
| | [Bmim]Cl, tetrabutylphosphonium chloride [$(C_4H_9)_4P$][Cl], and butylmethylpyrrolidinium chloride [$C_4$MPy][Cl] at 130°C for 10-24 h | Hoof, wool | Lovejoy et al. (*61*) |
| | 1-allyl-3-methylimidazolium [Amim]Cl, [Bmim]Cl, [Bmim]$NO_3$, and [Bmim]$HSO_4$ with $Na_2SO_3$ at 90-120 °C, ~90% dissolution | Feather | Ji et al. (*62*) |
| | 1-ethyl-3-methylimidazolium acetate [Emim][OAc] at 80°C, even dissolution achieved after 287 h, 12.7% yield after precipitation in ethanol | Bovine Hoof | Apostolidou. (*63*) |
| | 7-methyl-1,5,7-triazabicyclo(4.4.0)dec-5-ene levulinic acid [mTBDH][Lev] and [mTBDH][OAc] at 85°C for 3 h, then precipitated in water/ethanol, maximum dissolution capacity 13 wt% | Wool | Fang et al. (*20*) |
| Microwave irradiation | Microwave irradiation with 50-570 W power in superheated water at 150-180°C for 30-60 min, Mw around 3-8 kDa with loss of α-helix structures | Wool | Zoccola et al. (*64*) |
| | Microwave irradiation with max 1600 W power in superheated water at 180°C for 30 min, 31% yield, Mw around 5 kDa | Wool | Bertini et al. (*65*) |
| | Microwave irradiation with max 1200 W power of autoclave reactor at 160-200°C for 20 min, 71.83% yield of amino acids | Feather | Chen et al. (*66*) |
| Steam explosion | Saturated steam treatment at 220°C for 10 min, resulting in a dark-yellow slurry (solid residue and degraded keratin) with little α-helix remaining | Wool | Tonin et al. (*67*) |
| | Steam explosion at 1.4-2.0 MPa for 0.5-5 min, 65.78% yield with loss of secondary structures | Feather | Zhang et al. (*68*) |
| Thermal hydrolysis | Superheated water in a steel pressure cell at 220°C for 120 min, resulting in oligopeptides between 10-18 amino acids | Feather | Yin et al. (*69*) |

| | | | |
|---|---|---|---|
| | Superheated water in a thermal hydrolysis reactor at 100-220°C for 60 min, then filtration and evaporation. 40-70% yield of ~10 kDa amino acids | Dog hair | Tasaki (*70*) |
| Enzymatic/ microbial degradation | Incubation with esperase and 0.165 M L-cysteine in 20 mM borax buffer (pH 10.15) at 50°C for 15 h, over 95% product has less than 1 kDa Mw | Wool | Zhang et al. (*71*) |
| | Co-cultivation of B. licheniformis BBE11-1 and S. maltophilia BBE11-1, 81.8% degradation in 96 h, resulting in amino acids and soluble peptides | Feather | Peng et al. (*72*) |
| | Degradation with keratinase from soil metagenomes at 50°C for 48 h with 1% glutathione (pH 10), followed by dialysis for 3 days, 33.7% yield with 45/28 kDa Mw | Wool | Su et al. (*73*) |
| | Incubation with Bacillus sp. CN2 in fermentation medium at 37°C for 12-72 h, resulting in free amino acids and soluble peptides | Feather | Lai et al. (*74*) |
| Lithium bromide (LiBr) | Pre-treatment in HCl solution (pH 2) for 1 h, followed by extraction in 0.1 M LiBr, 0.5 M $NaHSO_3$, and 0.02 M SDS at 90°C for 4 h, dialyzed for 48 h, 50.2% yield | Wool | Zeng et al. (*75*) |
| | 8 M LiBr and 0.1 M DTT at 90°C for 36 h, followed by addition of NaCl after hot filtration, then stored at 4°C for 12 h to obtain keratin gel, 43.6% yield with shape memory | Wool | Cera et al. (*15*) |
| | 8 M LiBr and 0.1 M DTT at 90°C for 36 h, then acidified with HCl to pH 1.0-1.7, centrifuged to obtain subnatant gel, 49% total yield | Wool | Sun et al. (*19*) |
| | 0.3 M LiBr, 10-50 mg/ml L-cysteine, 0.5/1.0/1.5 M $Na_2S$/$Na_2S_2O_3$/$NaHSO_3$ at 60°C for 12 h, dialyzed for 72 h and dried. α-helix recovered, 10%-50% yield | Wool | Wang et al. (*76*) |

**Table S2. Summary of software and models used for simulation**

| Software/package | Purpose | Reference |
|---|---|---|
| NAMD 2.14 | Conduction of all molecular dynamics simulations | Phillips et al. (*40*) |
| VMD 1.9.3 | Visualization of results from molecular dynamics simulations | Humphrey et al. (*41*) |
| ColabFold 1.5.5 | Obtaining the initial atomistic protein structure | Mirdita et al. (*49*) |
| PyMOL 2.5.5 | N-terminus acetylation and C-terminus amidation of protein | - |
| PLUMED 2.8.3 | Examination of convergence of Metadynamics simulations | Tribello et al. (*50*) |
| tLEaP | Preparation of initial configurations of simulation box | Case et al. (*45*) |
| ff14SB | Force field for atomistic protein simulation | Maier et al. (*42*) |
| TIP3P | Water model for explicit solvation in molecular dynamics simulations | Price et al. (*43*) |
| Li/Merz | Ion parameters for monovalent ions in molecular dynamics simulations | Li et al. (*44*) |
| 2PT | Two-phase Thermodynamic (2PT) Model for calculation of water entropy | Lin et al. (*46*) |
| Colvars | Well-tempered metadynamics simulations for protein free energy landscape | Fiorin et al. (*37*) |
